# A single-cell atlas of the aging murine ovary

**DOI:** 10.1101/2023.04.29.538828

**Authors:** José V. V. Isola, Sarah R. Ocañas, Chase R. Hubbart, Sunghwan Ko, Samim Ali Mondal, Jessica D. Hense, Hannah N. C. Carter, Augusto Schneider, Susan Kovats, José Alberola-Ila, Willard M. Freeman, Michael B. Stout

**Affiliations:** Aging & Metabolism Research Program, Oklahoma Medical Research Foundation, Oklahoma City, OK, USA; Genes & Human Disease Research Program, Oklahoma Medical Research Foundation, Oklahoma City, OK, USA; Oklahoma City Veterans Affairs Medical Center, Oklahoma City, OK, USA; Nutrition College, Federal University of Pelotas, Pelotas, RS, Brazil; Arthritis & Clinical Immunology Research Program, Oklahoma Medical Research Foundation, Oklahoma City, OK, USA

**Keywords:** collagen, fibrosis, menopause, mouse, multinucleated giant cells, reproduction

## Abstract

Ovarian aging leads to diminished fertility, dysregulated endocrine signaling, and increased chronic disease burden. These effects begin to emerge long before follicular exhaustion. Around 35 years old, women experience a sharp decline in fertility, corresponding to declines in oocyte quality. Despite a growing body of work, the field lacks a comprehensive cellular map of the transcriptomic changes in the aging ovary to identify early drivers of ovarian decline. To fill this gap, we performed single-cell RNA sequencing on ovarian tissue from young (3-month-old) and reproductively aged (9-month-old) mice. Our analysis revealed a doubling of immune cells in the aged ovary, with lymphocyte proportions increasing the most, which was confirmed by flow cytometry. We also found an age-related downregulation of collagenase pathways in stromal fibroblasts, which corresponds to rises in ovarian fibrosis. Follicular cells displayed stress response, immunogenic, and fibrotic signaling pathway inductions with aging. This report raises provides critical insights into mechanisms responsible for ovarian aging phenotypes.

## INTRODUCTION

Ovarian aging has garnered significant attention in recent years due to a large proportion of women choosing to delay childbearing^1^, which often causes difficulty with conception and carrying a pregnancy to full-term^2^. As the ovary ages, the local microenvironment changes in ways that reduce oocyte quality and increases the rate of follicular depletion, which eventually results in menopause. Menopause is associated with accelerated systemic aging^3^, greater chronic disease burden^4–6^, and increased all-cause mortality risk^7^. Therefore, a deeper understanding of the mechanisms that underlie ovarian aging is critically important to extending female fertility and attenuating age-related chronic disease onset.

It is well-established that age-related ovarian follicular depletion is associated with increased mitochondrial dysfunction^8^, reactive oxygen species production^9,10^, inflammation^11–13^, and fibrosis^14,15^. However, very little is known about which cell-types develop these phenotypes first and/or dominantly contribute to the changing local microenvironment. Moreover, it remains unclear if cells in the follicle, stroma, or both play mechanistic roles in promoting follicular depletion and ovarian failure. Recent work has sought to unravel the potential role that ovarian stromal cells play in ovarian health and disease^12,15,16^. We and others have reported that markers of cellular senescence and fibrogenesis increase within the ovarian stroma with aging^11,12,15^, although the specific cell-types that become senescent and/or pro-fibrotic remain unknown. In addition to increased senescence and fibrotic markers, the ovarian stroma also accumulates multinucleated giant cells (MNGC) with advancing age, which may be related to the mechanisms that promote the aforementioned phenotypes^17^.

Due to the complex nature of ovarian function, which changes dynamically during aging, it has historically been challenging to elucidate the cell-type-specific mechanisms that promote follicular depletion and ovarian failure. Data showing age-related changes in ovarian transcriptional programs within cellular subtypes is limited, particularly within mice. Single-cell analyses of ovarian aging in non-human primates identified a downregulation of antioxidant programs in aged oocytes and increased apoptosis in aged granulosa cells (GC)^18^. Additionally, single cell analyses of human ovarian tissue is currently in progress^19^. However, mice represent the most utilized model organism for ovarian aging studies^20^ due to their short lifespan and ease of genetic manipulation for mechanistic studies. A spatially resolved analysis of murine ovaries made strides in identifying age-related changes in ovarian cell populations^21^. However, this study used reproductively senescent mice (15-month-old) and lacked the resolution to identify specific cellular subtypes and important immune populations. To add to this growing body of work, we performed single-cell RNA sequencing (scRNA-seq) to identify age-related transcriptional changes in a cell-type-specific manner within the murine ovary at 3 and 9 months of age. We chose to analyze 3- and 9-month-old mice because they remain reproductively active at both ages, yet this age interlude represents a period when follicular density decreases dramatically in conjunction with the emergence of age-related hallmarks^11^. This design and approach allowed us to determine how aging modulates cellularity and cellular phenotypes within the ovary prior to reproductive senescence, which is vital to developing pharmacological approaches for extending reproductive lifespan in females.

## RESULTS

### scRNA-seq of the adult murine ovary across reproductive ages

To evaluate age-related changes in the ovary, we collected ovaries from 3- and 9-month-old female mice (n=4/group) and performed scRNA-seq. The period from 3 to 9 months represents the time when the greatest decline in follicular reserve occurs and other hallmarks of aging begin to emerge^11^. Following quality control analyses, filtering, and doublet removal, 14,504 cells remained for characterization. Cellular distributions of the number of genes detected, number of molecules, and proportion of mitochondrial DNA before and after filtering are shown in Suppl. Fig. 1A. A mtDNA filtering parameter of 25% was used to ensure inclusion of oocytes because they are known to have more mitochondria than somatic cells^22,23^. Additionally, mitochondrial-dependent apoptosis is involved in follicular atresia^24^. Thus, a lower filtering threshold may result in loss of oocyte and follicular cells. Indeed, these clusters (CLUs) showed the highest mtDNA percentages among all identified CLUs (Suppl. Fig. 1B). Unbiased clustering and uniform manifold approximation and projection (UMAP) analysis revealed fifteen distinct cellular CLUs (Fig. 1A). One CLU was present in only one sample and was subsequently identified as oviduct contamination (Suppl. Fig. 1C), therefore these cells were removed from further analyses which left 14,349 total cells for downstream analyses. To assign cell-type identity, we used cell-type-specific markers previously reported in the literature.

**Fig. 1.**
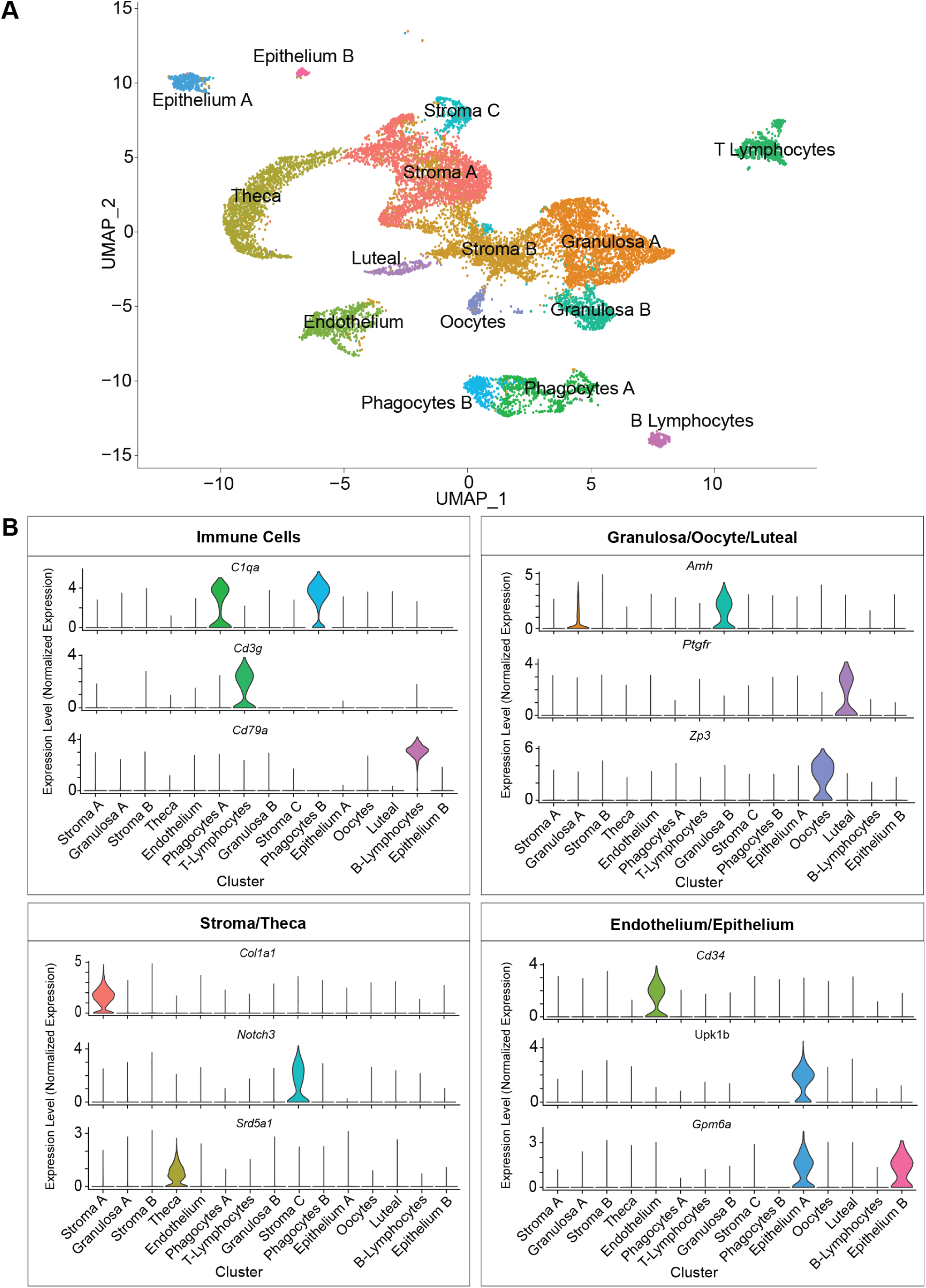
scRNA-seq of the murine ovary. Whole ovarian tissue was collected from 3- and 9-month-old C57BL/6J mice and processed for 10X Genomics 3’ scRNA-Seq. (A) UMAP plot of age-combined ovarian cells. Clustering analysis revealed 15 distinct ovarian cell populations. (B) Violin plots of specific marker genes for each ovarian cell type. scRNA-seq was performed in n=4 ovaries/age.

Collectively, the most common cell type was found to be stromal cells (N=5,671), which segregated into three CLUs. One stromal CLU, referred to as Stroma A, was characterized by having a major *Col1a1* transcriptional signature^25^, as well as other stromal markers. A second CLU, Stroma B, was identified by the expression of several stromal markers (*Bgn*, *Ogn*, *Dcn*, *Lum*, *Col1a1*^26^). However, this CLU did not have any exclusive markers that were highly enriched. A third stromal CLU, Stroma C, was characterized by *Notch3* expression (Fig. 1B), which is classically viewed as a pericyte marker^26,27^. The second most common most common cell type was found to be the GCs (N=3,334) which segregated into two distinct CLUs, both displaying high expression of *Amh*^18,28^ (Fig. 1B). Other CLUs were identified as theca cells (TC; N=1,637; *Srd5a1*^29^), phagocytes (two distinct CLUs; N=1,099; *C1qa*^30^), endothelial cells (N=798; *Cd34*^31^), T-lymphocytes (N=728; *Cd3g*^32^), epithelial cells (two distinct CLUs; N=450; *Upk1b*^33^ or *Gpm6a*^26^), oocytes (N=224; *Zp3*^18^), luteal cells (N=206; *Ptgfr*^34^), and B-lymphocytes (N=202; *Cd79a*^35^) (Fig. 1B). Feature plots displaying the specificity of these markers to each CLU can be found in Suppl. Fig. 1D.

Advancing age changed ovarian cellularity in our analyses (Fig. 2A-B, Suppl. Fig. 1E). The percentage of GCs was lower in the 9-month-old ovaries, which can be explained by declines in the number of primordial and tertiary follicles as well as a trending decline in secondary follicles observed in the aged mice (Fig. 2C-E). The most dramatic change in ovarian cellularity with advancing age was the greater than 2-fold increase in immune cells (Fig. 2A-B, Suppl. Fig. 1E).

**Fig. 2.**
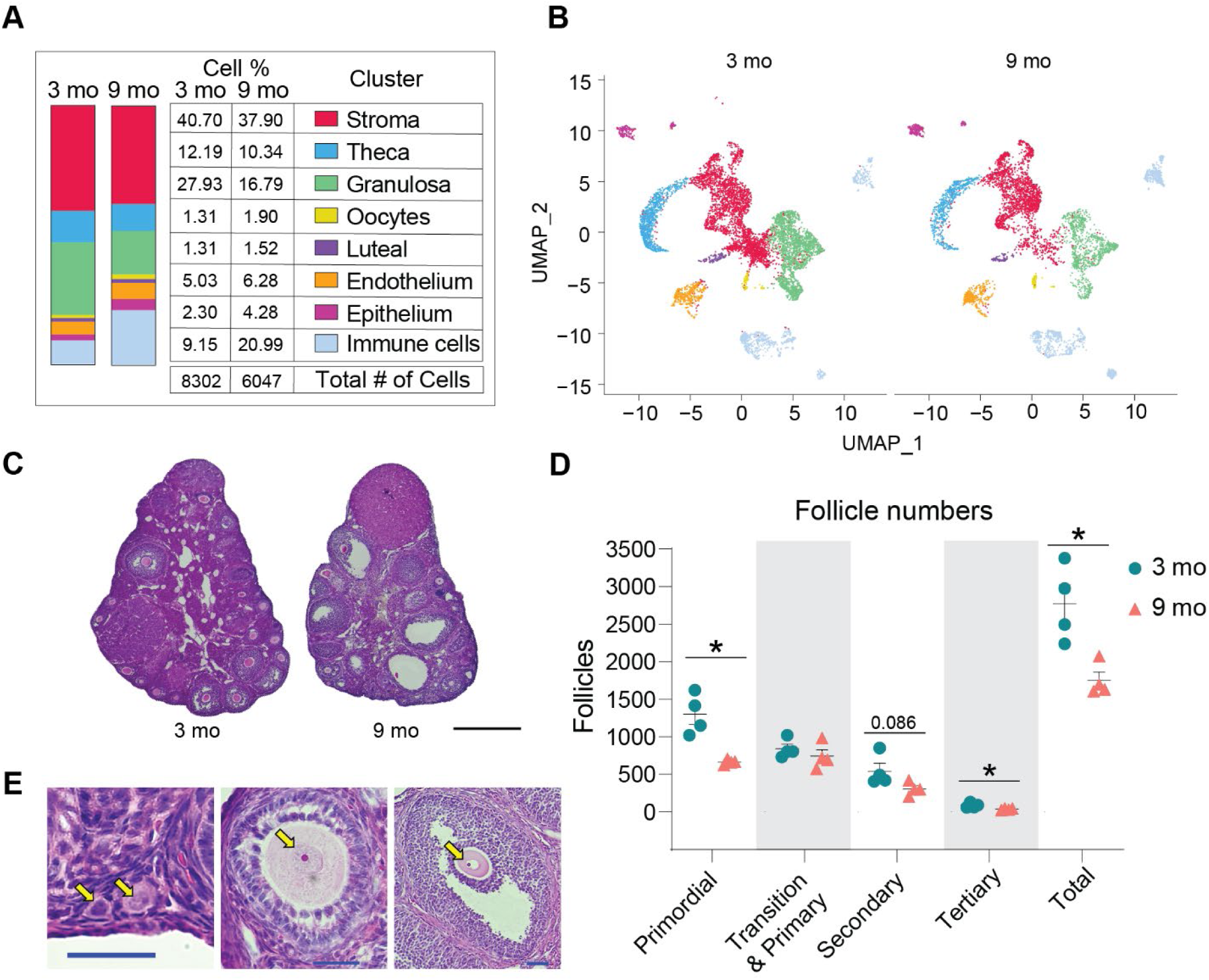
Age-related changes in ovarian cell populations. (A) The number and percentage of cells in broad categories of cell type identity by age. (B) UMAP plot of ovarian cells split by age. (C) Representative images of H&E stained ovaries. (D) Estimated number of ovarian follicles in 3- and 9-month-old ovaries. (E) Representative images of follicles stained by H&E. Arrows are pointing to a primordial, primary, secondary and tertiary follicles (from left to right). Data are presented as mean ± SEM. *, **, *** represent statistical difference (FDR<0.05, 0.01 and 0.005, respectively) by multiple two-tailed t-test with Benjamini, Krieger, and Yekutieli correction for multiple comparisons. Black scale bars represent 500µm and blue scale bars represent 50µm. scRNA-seq was performed in n=4 ovaries/age.

In this study, estrous cycle staging was not performed. However, we used a bioinformatic approach to determine which estrous cycle stage each mouse was in at the time of euthanasia. To do this, we initially integrated our data with the single-cell transcriptomic data from estrous-cycle-staged mice^26^. This report showed major differences in single-cell transcriptional outcomes within GCs across estrous cycle stages, with unique features being observed during proestrus and estrus. Following integration, we then generated UMAP plots of GCs from mice in each estrous cycle stage and observed unique CLUs during estrus and proestrus, which recapitulated the findings from Morris et al^26^. We then created UMAP plots of GCs from each biological replicate in our study to determine if they were in proestrus or estrus, which could potentially confound the interpretation of age-related transcriptional phenotypes (Suppl. Fig. 2A-D). We found that all 4 young mice, and 3 of the old mice, were in diestrus or metestrus, which are transcriptionally indistinguishable^26^. One old mouse, Sample 5, was found to be in proestrus. We then used the list of transcripts reported by Morris et al^26^. to be upregulated during proestrus within GCs to generate a module score for each sample, which enabled us to confirm that Sample 5 was is proestrus. This data is presented in violin plots (Suppl. Fig. 2E). We also performed a Principal Component Analysis (PCA) of differentially expressed genes with age in our samples and observed that the only distinguishable clustering is according to age (Suppl. Fig. 2F). Assessment of the potential impact of the proestrus sample on age-related phenotypic changes in GCs is discussed later in the manuscript following sub-cluster (SCL) analyses, although no major changes were observed.

To evaluate intercellular signaling networks, CellChat analyses were performed. The most notable changes in cell signaling occurred among the granulosa, stroma A, and epithelium CLUs with an overall increase in the number of interactions (Suppl. Fig. 3A) and interaction weights (Suppl. Fig. 3B) from 3 to 9 months-old. However, there was not a complete breakdown in cell-to-cell communication with aging.

### Ovarian immune cell changes with aging

After observing the increased proportion of immune cells within the aged ovary, we performed sub-clustering of immune cells to determine which specific populations were changed with aging (Fig. 3A). Upon re-clustering, the original immune cell CLUs separated into 13 SCLs (Fig. 3B). Cell type identification for each SCL was performed based on known gene expression profiles, and with the help of the Cellmarker2.0^36^ and ImmGen^37^ databases, with representative genes described in Fig. 3C. Since the overall immune cell proportion doubled from 3 to 9 months of age, percentages of total cells in each SCL were assessed and compared by age. Intriguingly, the cell types that showed the greatest increase with age were lymphoid, including B cells, conventional T cells, and innate-like T cells. The biggest difference in abundance with age was in a SCL containing Type 1 lymphoid cells, as defined by the expression of *Tbx21*^38^ and *Ifng*. These likely include CD8^+^ T cells, as well as other Type 1 innate-like T cells such as Natural Killer T Cells (NKT), Mucosal-associated invariant T cells (MAIT) and/or γδ T cells (γδT). Another SCL that significantly increased with age contained Type 17 lymphoid cells, as defined by the expression of *Rorc*^39^, which commonly represents innate lineage, and expression of *Zbtb16*^40,41^. Although transcriptional data cannot further discriminate within this SCL, it likely includes Type 17 NKTs, MAITs, γδTs and/or innate lymphoid cells (ILC) type 3, which was later confirmed by flow cytometry. Finally, the percentage of CD4^+^ T cells were also increased within the aged ovary (Fig. 3D). It should be noted that due to single cell transcriptomic limitations, other conventional T cells may also be contained in the CD4^+^ SCL. Conversely, neither natural killer (NK) nor ILC2s were significantly changed with age. Tissue macrophages and monocytes trended towards an increased proportion in the aged ovaries, but failed to reach statistical significance (Fig. 3D). One SCL was only present in the aged ovaries, which was identified as monocytes that express CD300e, which negatively regulates T-cell activation^42^. Hence, these CD300e^+^ monocytes might represent a compensatory mechanism to address the T-cell accumulation. However, this SCL might also represent blood monocyte contamination due to incomplete perfusion. Further investigation into this unique cellular population is warranted. Interestingly, although having an immune-like expression profile, two of the immune SCLs show no expression of *Ptprc*, the gene that codes for CD45, which is the most well-established immune cell marker. These two SCLs were identified as CD45^−^ immune-like cells A and B.

**Fig. 3.**
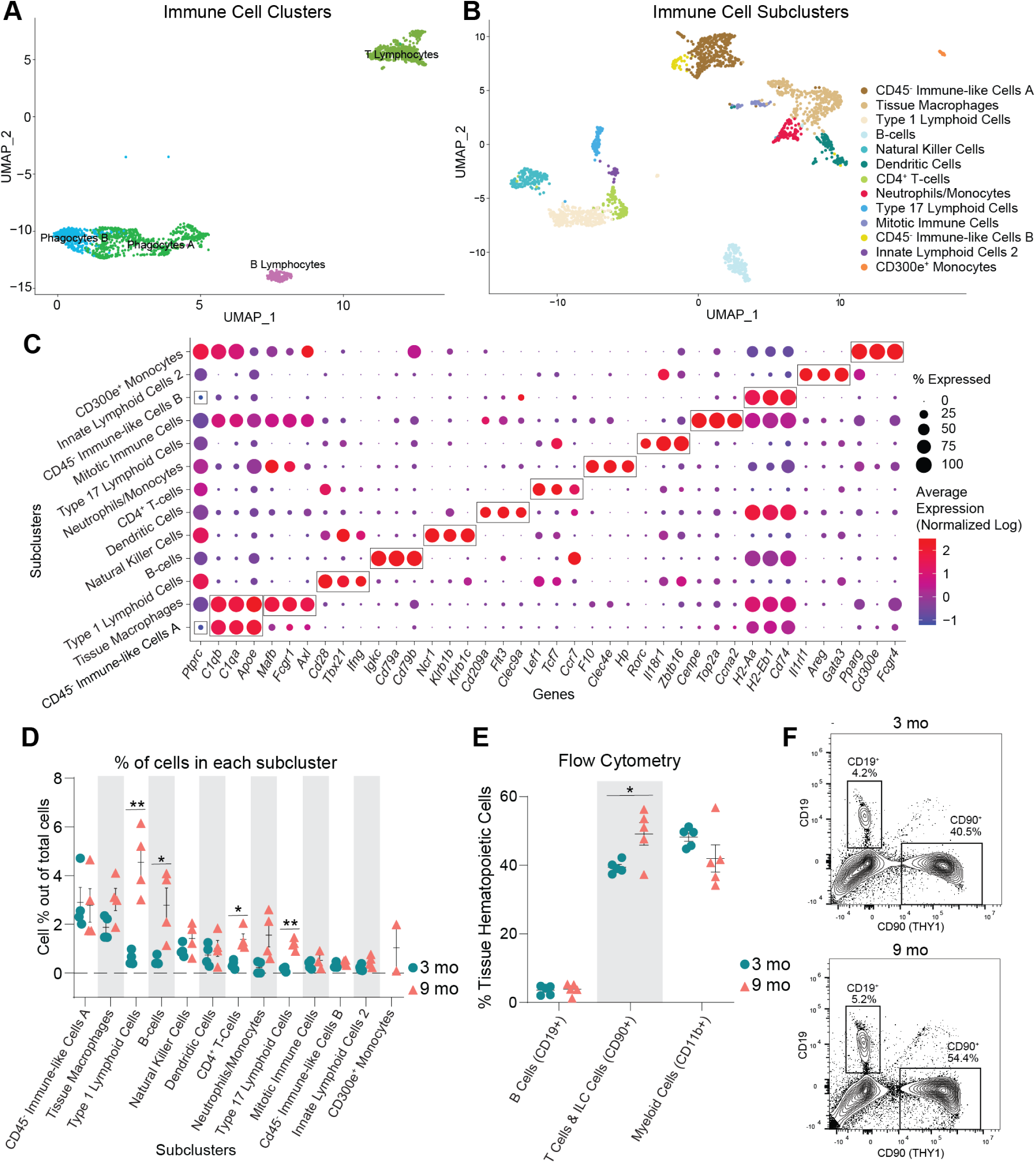
Immune cells accumulate in the ovaries with age. (A) UMAP of immune CLUs selected for sub-clustering analyses. (B) UMAP of immune SCL. (C) Dot plot of markers used for SCL cell-type identification. (D) Percentage of cells in each immune SCL out of total cells. (E) Percentages of immune cells out of tissue hematopoietic cells (after gating out intravascular hematopoietic cells) by flow cytometry. (F) Representative images of flow cytometry gating for CD19^+^ and CD90^+^ cells out of tissue hematopoietic cells (i.v. CD45^−^; CD45^+^) in young and aged ovaries (see Suppl. Fig 4A for the complete gating strategy). For flow cytometry, 6 ovaries from mice in the same phase of estrous cycle were pooled together to comprise each sample (n=5/age). scRNA-seq was performed in n=4 ovaries/age. Data are presented as mean ± SEM. *, **, *** represent statistical difference (FDR<0.05, 0.01 and 0.005, respectively) by multiple two-tailed t-test with Benjamini, Krieger, and Yekutieli correction for multiple comparisons.

To confirm our transcriptional findings related to ovarian immune cell accumulation, we performed high dimensional flow cytometry to identify age-related differences in the fraction of distinct immune cell types within the ovary. We first assessed the percentage and total number of broadly defined immune cell types, including B cells (CD19^+^), T cells & ILCs (CD90^+^), and myeloid cells (CD11b^+^). We observed a higher percentage and number of CD90^+^ cells (T cells and ILCs) in aged ovaries (Fig. 3E-F, Suppl. Fig. 4A), which is aligned with the scRNA-seq findings. However, we did not observe changes in the percentage or number of myeloid cells or B cells (Fig. 3E-F, Suppl. Fig. 4A). Although the lack of change in myeloid cells was consistent with the scRNA-seq data, the B cells were incongruent. The proportion of B cells were found to increase with aging by scRNA-seq analyses, although this was not confirmed by flow cytometry. UMAP visualization of the flow cytometry data showed major shifts in immune cell subpopulations within the aging ovary (Fig. 4A). We then further characterized the CD90^+^ population and observed an increase in the percentage of CD8^+^ T cells, double negative αβ T cells (DN), MAITs, NKTs, and γδTs and a decrease in CD4^+^ αβ T cells and ILCs in the aged ovaries (Fig. 4B, Suppl. Fig. 4B). To compare and contrast these findings with the scRNA-seq clustering, we analyzed the CD90^+^ cells based on expression of the effector-type-determining transcription factors for Type 17 lymphoid cells (RORγT) and Type 1 cells (TBET) (Suppl. Fig. 4C). The increase in Type 1 lymphoid cells observed by scRNA-seq was confirmed by flow cytometry and further determined that the increase was driven by Type 1 NKTs, while Type 1 DN and γδTs proportions decreased with age (Fig. 4C-D). All Type 17 lymphoid cell subpopulations increased in proportion with age (DN, NKT, γδT), although the greatest magnitude of change occurred in Type 17 γδTs (Fig. 4C-D). Although the scRNA-seq data suggested the presence of ILC2, but not ILC1 subpopulations, the flow cytometry data identified both ILC1s (TBET^+^) and ILC2s (GATA3^hi^). The flow cytometry data indicated a decrease in ILC1 percentages with advancing age (Fig. 4E-F). To determine the potential impact of estrous cycle stage on age-related changes in ovarian immune cell populations, we performed vaginal cytology on mice prior to euthanasia and ovarian flow cytometry analyses. We then plotted the mean fluorescence intensity of all markers assessed in each ovarian sample in addition to identifying which estrous cycle stage each mouse was in. We found that samples clustered by age, regardless of estrous cycle stage (Suppl. Fig. 4D), thereby indicating that estrous cycle stage is not a primary contributor to age-related changes in immune cell accumulation.

**Fig. 4.**
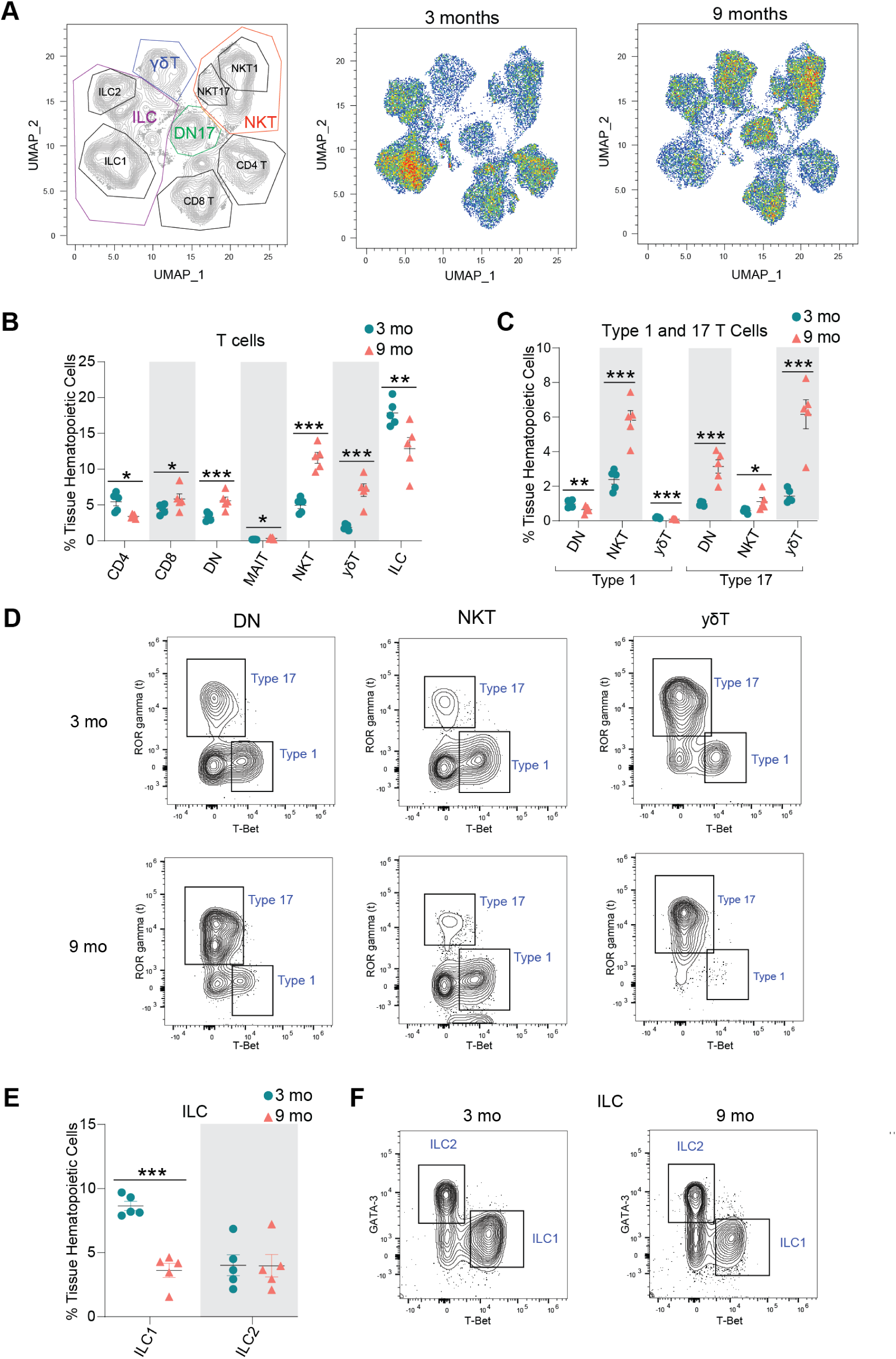
Lymphocyte populations are altered in the aged ovary. (A) UMAP of flow cytometry panel of lymphoid cells, annotated using conventional gating strategies. The number of cells plotted was normalized to better observe differences in distribution of the populations. (B) Percentage of T cell subpopulations out of tissue hematopoietic cells. (C) Percentage of Type 1 and Type 17 T cells out of tissue hematopoietic cells. (D) Representative gating of Type 1 and Type 17 lymphoid cells in young and aged ovaries. (E) Percentage of ICL1 and ILC2 out of tissue hematopoietic cells. (F) Representative gating of ILCs in young and aged ovaries. For flow cytometry, 6 ovaries from mice in the same phase of estrous cycle were pooled together to comprise each sample (n=5/age). scRNA-seq was performed in n=4 ovaries/age. Data are presented as mean ± SEM. *, **, *** represent statistical difference (FDR<0.05, 0.01 and 0.005, respectively) by multiple two-tailed t-test with Benjamini, Krieger, and Yekutieli correction for multiple comparisons.

In addition to an overall increase in immune cells within the aged ovary, previous reports have also noted the accumulation of MNGCs as a hallmark of ovarian aging^17,43^. Although MNGCs were certainly removed from our single-cell suspensions during filtration, we were able to observe them in 9-month-old ovaries via histological assessments. Similar to prior reports^11,17,44^, we observed MNGCs in the aged ovary (Suppl. Fig. 5A) and the presence of these structures corresponded to a rise in lipofuscin positivity (Suppl. Fig. 5B), which has been reported to be a marker of cellular senescence^45^. However, it should be noted that lipofuscin positivity strongly associates with increased numbers and size of MNGCs, which may not actually be cellular senescence as classically defined. Given the increased number of immune cells in the aged ovary that inherently express senescence-related genes (Suppl. Fig. 6A-B), it appears likely that ‘ovarian cellular senescence’ prior to estropause may represent increased immune cell accumulation. However, it should be noted that *Cdkn1a* and *Cdkn2d* expression increased in Type 17 lymphoid cells (Suppl. Fig. 6B-D) with advancing age, which could potentially represent a small subpopulation of cells that enter cellular senescence in the ovary.

### Sub-clustering of stroma and theca cells reveals age-related phenotypic changes

Several changes in the stroma have been observed with ovarian aging and are apparent prior to follicular exhaustion, including cellular senescent signatures^11^ and collagen deposition^12,14,15^. To identify potential cellular and molecular mediators of these phenotypes, we performed sub-clustering of the stroma and TC CLUs (Fig. 5A-B). TCs were included with stroma since they dynamically differentiate from fibroblastic stromal cells during follicular maturation, and thus, share similarities with stromal cells^36^. This sub-clustering resulted in six distinct SCLs (Fig. 5B). Enriched markers in each SCL were used to infer cell-type-specificity (Fig. 5C). Notably, there was a SCL that expressed *Inhba*^37,38^, a marker of GCs, which was distinct, and thus, easily separated from other stroma and TC SCLs. The smooth muscle SCL was identified by the expression of *Myh11*^46,47^, *Actg2*^26^ and *Cnn1*^48^. The pericyte SCL was enriched for *Notch3*, *Rgs5* and *Ebf1*^26^. The fibroblast-like SCL had enrichment of *Dcn, Mgp*^40^, and *Lum*^26^. Whereas, the TC SCL presented high expression of steroidogenic genes (*Cyp11a1*, *Cyp17a1*^36^, and *Mgarp*^26^). The early TC SCL also presented high levels of steroidogenic genes such as *Ptch1* and *Hhip*^41^ in addition to high expression of a fibroblast gene (*Enpp2*^42^), suggesting that these cells are in transition from stromal fibroblast-like cells to TCs.

**Fig. 5.**
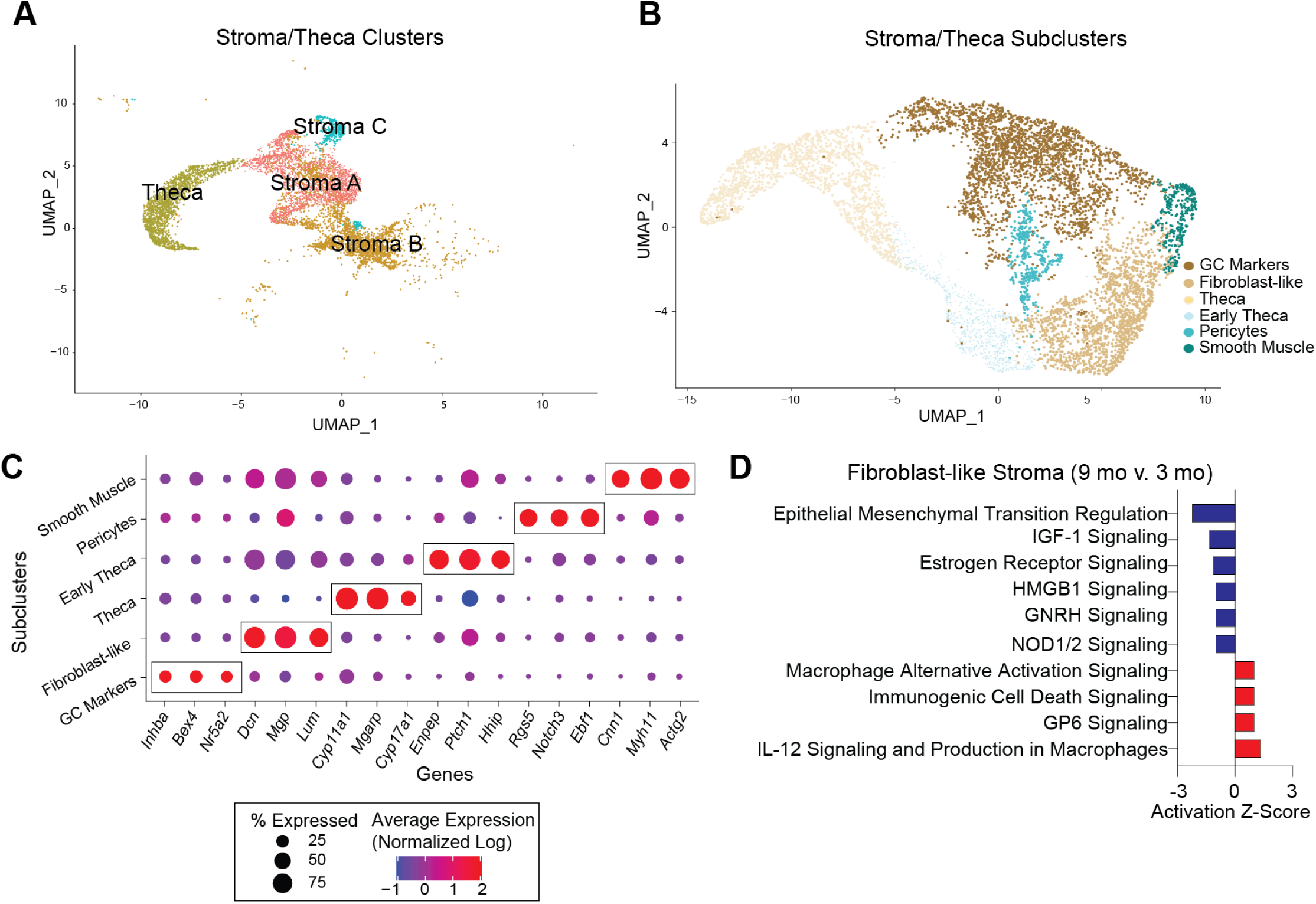
Sub-clustering of stroma and theca cells. (A) UMAP of stroma and TC CLUs used for sub-clustering (B) UMAP of stroma and TC SCLs. (C) Dot plot of markers used for SCL cell-type identification. (D) IPA canonical pathways indicating activation or inhibition of specific pathways by aging in stromal fibroblast-like cells. scRNA-seq was performed in n=4 ovaries/age.

After SCL identification, a list of genes differentially expressed by the different ages in each SCL was imported into the IPA software Ingenuity Pathway Analysis (IPA) to infer biological function. Fibroblast-like stromal cells showed an age-related upregulation of pathways related to tissue remodeling^49,50^ (Fig. 5D). Tissue remodeling occurs after ovulation and in cases of follicular and luteal atresia^16^. To our surprise, collagen expression was unchanged by aging in ovarian fibroblast-like SCL (Fig. 6A), despite greater collagen deposition by 9 months of age (Fig. 6B). Interestingly, the collagenase pathway was downregulated in the fibroblast-like SCL (Fig. 6C). Further analyses revealed that the expression of matrix metalloproteinase 2 (*Mmp2*), a key enzyme involved in collagen degradation^51^, was decreased in the fibroblast-like SCL with advancing age (Fig. 6D). We also observed an age-related decrease in MMP2 protein in the ovarian stroma by immunofluorescence (Fig. 6E-F). These findings suggest that ovarian collagen accumulation is at least partially mediated by a reduction in collagen degradation, which supports our transcriptional findings. Fibroblast-like cells also displayed a downregulation in hormonal signaling (estrogen receptor and GNRH) (Fig. 5D), which is somewhat surprising given that hormone levels, estrous cyclicity, and fertility are generally stable at 9 months of age^52^. These results indicate that changes in the local microenvironment may contribute to endocrine dysfunction in the ovarian stroma independent of changes within the follicle. In contrast to the fibroblast-like cells, TCs showed a significant upregulation in several upstream regulators of fibrogenesis including TGFB1, TGFB2, and SMAD3 (Fig. 6C), which suggests TCs may be one of the earliest cell-types involved in generating the signaling cascade for collagen production and deposition. When looking at TGF-β intercellular communication among the original CLUs, we note that TCs signal to several cell-types in the young ovary (Fig. 6G), which was expected, due to its important role in intrafollicular cell-to-cell communication^53^. In contrast, TGF-β signaling in TCs from aged ovaries becomes exclusive to immune cells (Fig. 6H). TGF-β signaling has previously been associated with the regulation of chemotaxis, activation, and survival of lymphocytes^54,55^, which may explain our finding of increased lymphocyte accumulation in the aged ovary. Additionally, the Granulosa B CLU only displayed TGF-β signaling in the old ovary (Fig. 6G-H). These data suggest that interactions between ovarian follicular cells and immune cells are likely important for perpetuating fibrotic signaling with aging. TCs also displayed a modest age-related upregulation in inflammation and cell stress response pathways (Fig. 6I), which were mirrored by upstream regulators involved in cellular proliferation including MTOR, YAP1, and RB1 (Fig. 6J).

**Fig 6:**
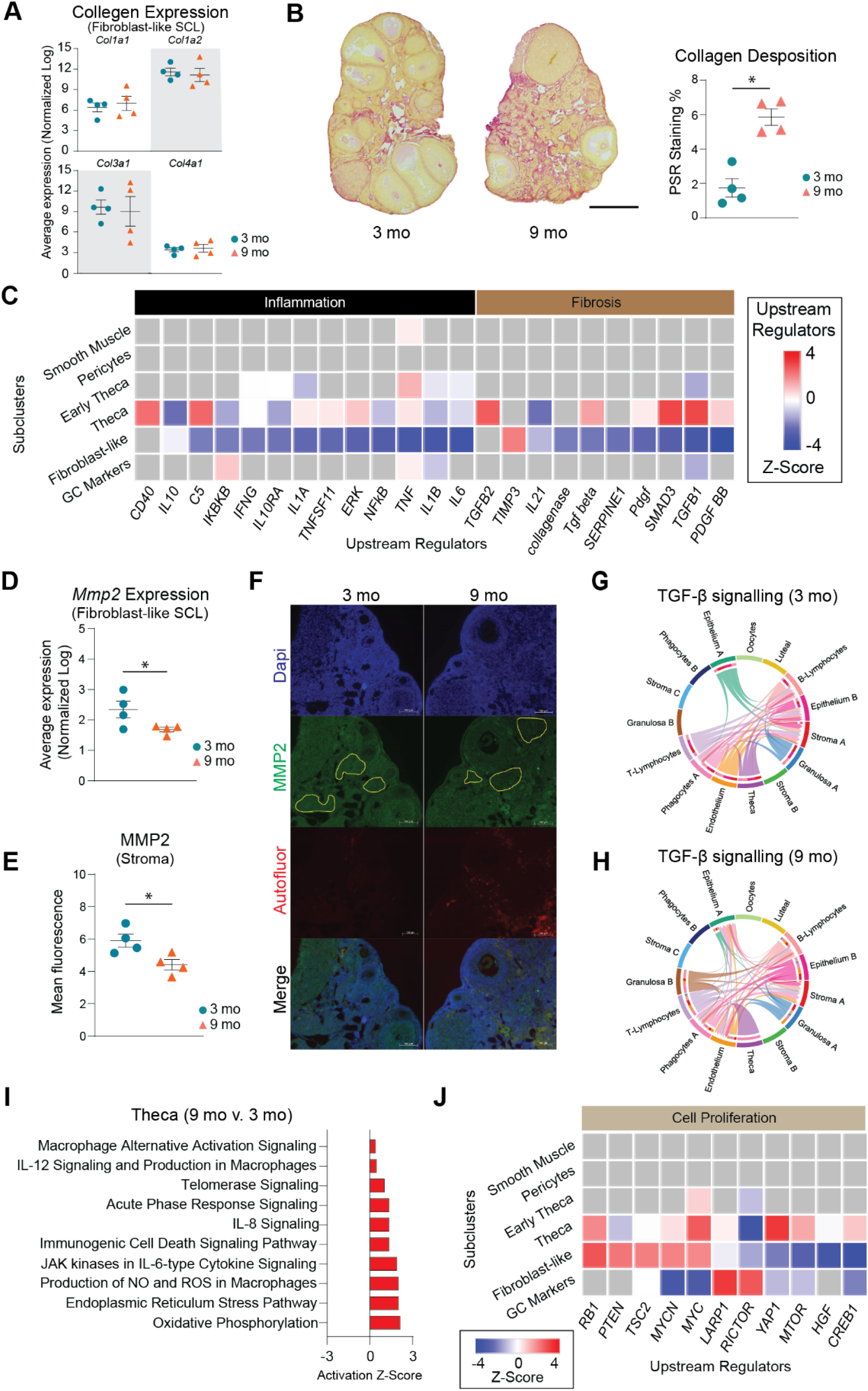
Biological significance of altered pathways in ovarian stroma and theca SCLs. (A) Expression of collagen genes in the fibroblast-like stroma SCL does not change with age. (B) PSR staining of collagen deposition in 3- and 9-month-old ovaries. (C) IPA upstream regulator analyses of age-related changes in stroma and TC SCLs (9 months-old vs. 3 months-old, z-score) related to inflammation and fibrosis. (D) Expression of Mmp2 gene in the fibroblast-like stroma decreases with age. (E) MMP2 protein is decreased in aged ovarian stroma by immunofluorescence. (F) Representative images of immunofluorescence of MMP2 (green), DAPI (blue) and autofluorescence (red). Yellow lines show representation of stromal regions analyzed, avoiding follicles and auto-fluorescent regions. (G-H) CellChat chord diagrams of the TGF-β signaling pathway interactions in the (G) 3-mo and (H) 9-mo ovarian CLUs. (I) IPA canonical pathways indicating activation of specific pathways by aging in TCs. (J) IPA upstream regulator analyses of age-related changes in stroma and TC SCLs (9 months-old vs. 3 months-old, z-score) related to cell proliferation. (n=5/age). scRNA-seq was performed in n=4 ovaries/age. Data are presented as mean ± SEM. *, **, *** represent statistical difference (p<0.05, 0.01 and 0.005, respectively) by one-tailed t-test (A-B) or two-tailed t test (D-E). Black scale bars represent 500µm and white scale bars represent 100µm.

### Sub-clustering of granulosa, oocytes and luteal cells reveal age-related pro-inflammatory changes

Cellular populations of GCs, oocytes, and luteal cells were sub-clustered into 6 distinct SCLs (Fig. 7A-C). GCs were further segregated into four distinct SCL that were identified as being part of follicles that were in different stages of development. The SCLs included preantral, antral, mitotic, and atretic GCs. GCs signal to oocytes to provide cues related to the local ovarian microenvironment, in addition to converting androgens to estrogens which are released into the systemic circulation for feedback signaling in the brain^56^. GCs from immature follicles that have not yet developed an antrum are referred to as preantral GCs and were identified by enrichment in *Igfbp5*^57^ and *Gatm*^58^ expression. GCs from antral follicles (antral GCs) were identified by enrichment of *Inhbb*^59,60^. A separate GC SCL, which we refer to as mitotic GCs, expressed *Inhbb* similarly to antral GCs, but displayed enriched expression of cell division markers *Top2a*^61^ and *Racgap1*^62^. Lastly, GCs from follicles undergoing atresia were found to be enriched for *Pik31p1*^26,63^, *Itih5*^26^ and *Ghr*^26^. Oocytes were easily identified by the expression of classical markers *Gdf9*, *Ooep* and *Zp3*^18^. Luteal cells, which consist of remnant GCs and TCs following ovulation and constitute the corpus luteum^64^, were identified by the expression of *Ptgfr*^34^.

**Fig. 7:**
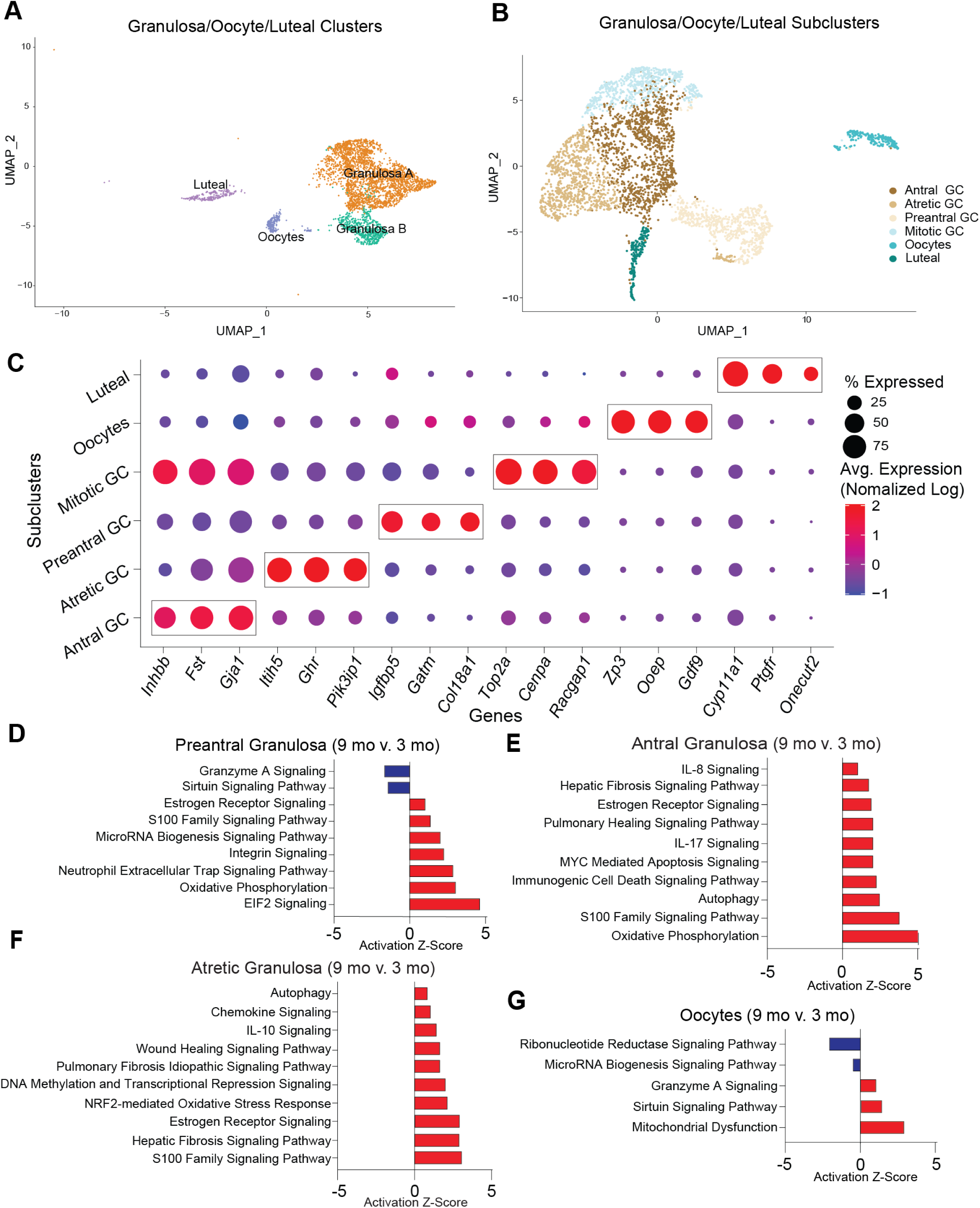
Sub-clustering of granulosa cells, oocytes and luteal cells. (A) UMAP of granulosa, oocytes and luteal cells CLUs used for sub-clustering (B) UMAP of granulosa/oocytes/luteal SCLs. (C) Dot plot of markers used for SCL cell-type identification. (D-G) Z-scores indicating activation or inhibition of pathways during aging by IPA analysis in (D) preantral GC, (E) antral GC, (F) atretic GC, and (G) oocytes. scRNA-seq was performed in n=4 ovaries/age.

As alluded to above, differentially expressed genes from each SCL were examined with IPA software to infer biological function. Interestingly, preantral, antral, and atretic GCs all displayed an age-related increase in pathways related to pro-inflammatory stress and fibrogenesis (Fig. 7D-F). These observations corresponded to a significant increase in the mitochondrial dysfunction pathway within oocytes (Fig. 7G). Upstream regulators related to pro-inflammatory stress that were found to be increased in GC SCL included IFNG, TNF, IL15, and IL1A (Suppl. Fig. 7A). Additionally, upstream regulators related to fibrogenesis were found to be increased in GC SCL including TGFB1, Tgf beta, and SMAD3 (Suppl. Fig. 7A). As expected, the atretic GC SCL was enriched for changes in pro-inflammatory stress and fibrosis pathways, with 9-month-old mice having greater activation of these pathways than 3-month-old mice. Therefore, it appears that aging exacerbates inflammatory and fibrotic responses that are required for ovarian remodeling following ovulation and follicular atresia.

We also examined intercellular C-C motif chemokine ligand (CCL) signaling due to its role in immune cell migration and chemotaxis. We found that by 9 months of age the granulosa CLUs expressed CCL signaling molecules, which was not observed in 3-month-old mice (Suppl. Fig. 7B-C). At 9 months of age, the GC CCL signaling was exclusive to the phagocyte A CLU. This suggests that inflammatory signaling from the follicle may contribute to alterations in immune cell phenotypes as a function of age. Interestingly, there was a breakdown in the anti-Mϋllerian hormone (AMH) signaling pathway between oocytes and other follicular cells by 9 months of age (Suppl. Fig. 7D-E). The AMH pathway is critical for regulation of follicular maturation^65^ and is commonly used as a marker of follicular reserve^66^. Thus, this breakdown in signaling signifies early changes that may impact reproductive outcomes.

Since GC transcriptional changes have been reported during proestrus, coupled with one of the 9-month-old mice (Sample 5) analyzed herein was in proestrus, we performed additional analyses to ensure that estrous cycle stage did not confound our age-related interpretations. For this analysis, we removed the genes that were found to be differentially expressed in both proestrus^26^ and aging in granulosa SCLs, and reran the pathway analyses. The removal of these genes did not affect the age-related changes in GCs. As a secondary confirmation, we then completely removed Sample 5, reran the pathway analysis again, and found that age-related outcomes were unchanged (Suppl. Fig. 7F). These observations provide additional evidence that our interpretations related to the mechanisms that promote ovarian aging are unaffected by estrous cycle stage.

### Sub-clustering of endothelial and epithelial cells reveals only minor age-related changes

Endothelial cells are essential components of blood and lymphatic vessels. The ovary is a highly vascularized organ with blood vessels that travel through connective tissue to assist with hormone/nutrient trafficking and waste removal^67^. The ovary also has a rich lymphatic network closely associated with the blood vasculature that is involved in immune cell trafficking^68^. Epithelial cells comprise the ovarian surface, facilitate repair following ovulation, and dynamically expand and contract during cyclic ovarian changes^69^. In addition, epithelium cells share a common progenitor with GC^70^. Endothelium and epithelium CLUs were sub-clustered (Fig. 8A) and resulted in four distinct SCLs (Fig. 8B). Vascular and lymphatic endothelium were separated and identified by the enrichment of specific cellular markers (Fig. 8C). Vascular endothelial cells showed enrichment of *Flt1*^71^ and *Mmrn2*^72^, whereas lymphatic endothelial cells were enriched for *Lyve1*^73^ expression.

**Fig. 8:**
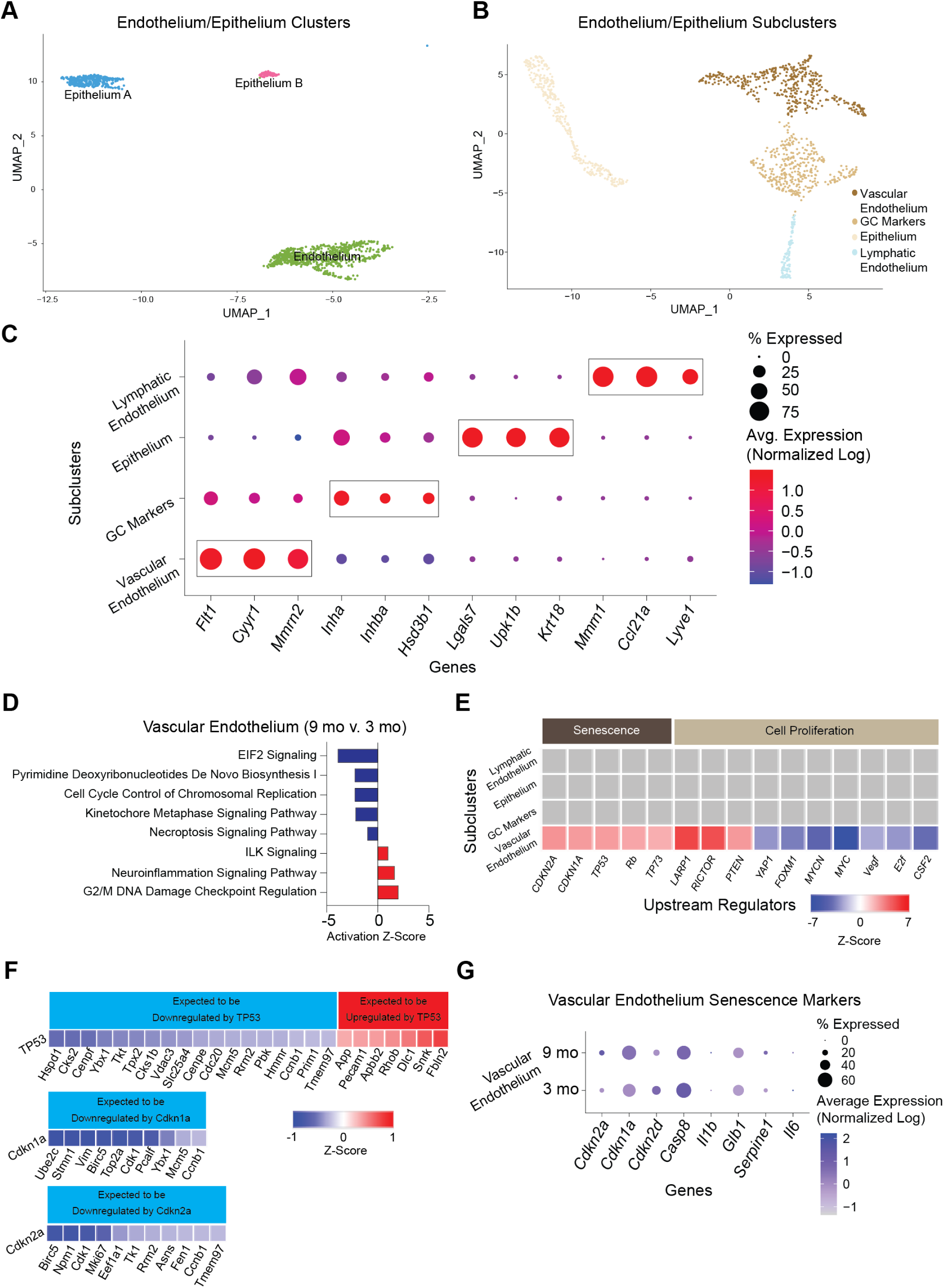
Sub-clustering of endothelial and epithelial cells. (A) UMAP of endothelium and epithelium CLUs used for sub-clustering. (B) UMAP of endothelium/epithelium SCLs. (C) Dot plot of markers used for SCL cell-type identification. (D) Z-scores indicating activation or inhibition of pathways altered with aging by IPA analysis in vascular endothelial cells. (E) Z-scores indicating activation or inhibition of upstream regulators during aging by IPA analysis in endothelium/epithelium SCLs. (F) Z-scores indicating up or downregulation of genes by TP35, Cdkn1a and Cdkn2a genes. (G) Dot plot of expression of cell senescence markers in vascular endothelial cells in 3- and 9-month-old ovaries. scRNA-seq was performed in n=4 ovaries/age.

Similar to findings in the stroma/theca SCLs, one of the endothelium/epithelium SCLs was found to be enriched for GC markers and was removed from further analyses. No age differences were observed in this GC SCL. The only epithelium/endothelium SCL that displayed age-related changes in the pathway analyses was the vascular endothelium. This SCL was found to have a modest increase DNA damage regulation (Fig. 8D). Another interesting observation was that upstream regulators of cellular senescence (TP53, CDKN1A, CDKN2A) were found to be increased with aging in vascular endothelial cells (Fig. 8E). A further analysis of genes that are downstream of TP53, CDKN1A, and CDKN2A were found to be altered in directionality consistent with the activation of these upstream regulators (Fig. 8F). This is consistent with literature that indicates vascular endothelial cells are highly susceptive to becoming senescent^74^. In the ovary, cyclicity requires constant vascular remodeling^67^, which we speculate may promote senescence in endothelial cells. However, despite observing increased TP53, CDKN1A, and CDKN2A upstream regulator activity in the vascular endothelium SCL, the expression of *Cdkn1a* and *Cdkn2a* was unaffected by aging (Fig. 8G), whereas *Tp53* was not detected in our scRNA-seq analysis. Therefore, it remains unclear if the vascular endothelium is a source of senescent cells in the aged ovary.

## DISCUSSION

In this report, we assessed age-related changes in the murine ovarian transcriptome at single-cell resolution. During aging, ovarian follicular reserve declines, and there is a concomitant deterioration of the ovarian microenvironment as evidenced by increased inflammation and fibrosis. Importantly, these changes occur prior to follicular exhaustion^11^ and likely contribute to decreased oocyte quality and diminished reproductive success until ovarian insufficiency occurs. However, specific cellular contributions to ovarian aging phenotypes are not yet elucidated, limiting the development of interventional approaches to extend female fertility. Whole ovarian bulk transcriptomic assessments can provide results that are difficult to interpret due to potential changes in cell heterogeneity that occur during aging. Cell sorting can overcome this limitation but relies on specific antibodies or cre-reporter systems to target specific cell-types. scRNA-seq, on the other hand, is a useful tool to simultaneously measure transcriptomic changes of all ovarian cell types during the aging process. Previous ovarian scRNA-seq have identified the molecular signatures of specific cell-types and changes in individual cell populations^18,26,28,58,75,76^. In mice, estrous cycle^26^ and primordial follicle assembly^75^ result in dynamic changes in specific ovarian cellular populations. However, the specific cellular changes occurring during ovarian aging are still being elucidated, especially with regard to the critical period of diminished fertility that occurs long before follicular exhaustion. Since mice are the primary model organisms used for ovarian aging experiments^20^, the single-cell ovarian aging atlas presented herein serves as a crucial resource for the field. In non-human primates, scRNA-seq provided mechanistic insights into changes associated with ovarian aging^18^. However, this study was conducted with animals in the peri-menopausal state^77^, when alterations in cyclicity and hormone levels are perturbed. Our primary goal was to identify early changes that occur in reproductively aged mice prior to the peri-estropausal period. At 9 months of age, mice experience declined ovarian reserve^11^, but remain fertile^78^, modeling a critical age when women seek to have children and experience difficulty conceiving^2^. A marked increase in embryonic aneuploidy is observed in embryos from women starting around 35 years of age^79^, suggesting a loss in oocyte quality. With our approach, we differentiated all of the main cell-types expected in the ovaries and further subdivided them to determine specific cellular populations altered by the aging process.

Our scRNA-seq results show that the proportion of immune cells in the ovary doubles between 3 and 9 months of age. The most dramatic increase in immune cells within the aged ovaries occurred in Type 1 and Type 17 lymphoid cells. A recent scRNA-seq assessment of ovarian immune cells reported an increase in CD4^−^CD8^−^ T cells, deemed “double negative T cells”, in aged ovaries^80^. Many of the Type 1 and Type 17 lymphoid cells described in the present study, including αβ DNs, Type 17 NKTs, MAITs, γδTs and a significant proportion of Type 1 NKTs, are CD4^−^CD8^−^ and would therefore be included within the broad “double negative T cell” population described in the previous study. Consistent with the prior report, our data indicates that many of these populations increase with age. However, our data provides better granularity and revealed specific populations of Type 1 and Type 17 lymphoid cells changing in the aging ovary that had not previously been evaluated.

Specifically, we determined that the immune subpopulations that most consistently increased in the aged ovary were Type 1 NKTs and Type 17 γδTs, both of which are innate T cells. Thus, our data raise interesting questions about the functional roles of innate T lymphocytes in both young and old ovaries. Innate T cells are tissue resident cells thought to contribute to the maintenance of tissue homeostasis and serve as sentinel cells to detect infectious agents, neoplastic cells, or aberrant cellular damage^81^. In the female reproductive tract, innate lymphocytes display high functional diversity with both beneficial and detrimental effects^82^. However, the role of innate lymphocytes in the ovary is an unexplored subject. Type 17 γδTs play critical roles during fibrosis, in a variety of organs, such as liver^83,84^, kidney^85^, lung^86^ and heart^87^. However, whether these cells are protective or deleterious in fibrosis appears to be tissue- and context-dependent^88,89^. For instance, age-related accumulation of Type 17 γδTs contributes to chronic inflammation in adipose tissue^90^. In the ovary, expansion of the Type 17 γδTs could either promote fibrosis or be a compensatory attempt to dampen the fibrotic environment. Little is known about the role of Type 1 NKTs in the ovary. In general, NKTs provide important immune responses during injury, repair, inflammation, and fibrosis in the liver and lung^87^. Further mechanistic studies assessing the roles of these cells in ovarian fibrosis and inflammation are warranted.

Under normal conditions, tissue resident lymphocytes are maintained through slow homeostatic proliferation^91^, as opposed to adaptive T cells which are generally recruited from circulation^92^. The mechanisms underlying increased innate lymphocyte numbers in aged ovaries may be a result of increased proliferation in response to tissue remodeling and/or local inflammatory cytokine production. This hypothesis is consistent with increased IFNG, TNF, IL15, and IL1A upstream regulator activity in GC SCLs. Alternatively, circulating lymphocytes could be recruited into tissue in response to similar pro-inflammatory signals^93^. Chronic ovarian inflammation caused by autoimmune disease also promotes lymphocyte accumulation^94^, and women with autoimmune diseases often experience ovarian insufficiency^95^. This suggests an association between the accumulation of lymphocytes and age-related follicular declines, which merits future interrogation.

Although we detected an increase in the proportion of B cells within the scRNA-seq data at 9 months of age, this was not confirmed by flow cytometry. A previous report showed that the proportion of ovarian B cells slightly increased from 2 to 6 months of age, but then returned to basal levels by 12 months of age^13^. This suggests that B cell numbers are highly variable in the aging ovary. Within our scRNA-seq data, we also detected an increase in CD4^+^ T cells in the 9-month-old ovary. Although the proportion of CD4^+^ T cells decreased between 3 and 9 months of age by flow cytometry, the absolute number of these cells increased with age. The accumulation of CD4^+^ T cells was previously reported in the aged ovary^13^, although these changes were not observed until 12 months of age. This suggests that the accumulation of CD4^+^ T cells may begin to occur at 9 months of age.

The literature addressing age-related changes in ovarian myeloid cells is conflicting^13,80,96^, which may be due to differences in the methods employed and ages evaluated. We found no significant changes in ovarian myeloid cell accumulation by 9 months of age through scRNA-seq or flow cytometry analyses. At this juncture, it remains unclear if myeloid cells actually increase in the ovary with advancing age, but the fusion of monocytes and macrophages into MNGCs^97^ could confound these interpretations. As addressed earlier, MNGC accumulation is a hallmark of the aged ovary^49^. Previous reports suggest that MNGCs may contribute to the age-related deterioration of the ovarian microenvironment^17^. However, the mechanisms that promote MNGC formation and their physiological ramifications in the ovary remain unresolved. The most well-characterized MNGCs are osteoclasts, which are implicated in bone remodeling through phagocytosis of apoptotic cells and collagen fragments^98^. In the ovary, MNGCs may also serve similar functions because phagocytic processes control tissue remodeling during follicular atresia, corpora lutea regression, and ovulation. Interestingly, estrogen induces osteoclast apoptosis, therefore, menopausal-related loss of estrogen action accelerates bone loss in women^99^. Similarly, decreased estrogen signaling in the aged ovary may promote MNGC accumulation. Lymphocytes, which are increased in the aged ovary, also contribute to MNGC formation through cytokine-mediated mechanisms^100^. For example, IL-17A (the primary cytokine produced by Type 17 lymphoid cells) is associated with osteoclast formation^101^, which suggests that Type 17 lymphoid cells may contribute to MNGC formation in the aged ovary. Similarly, IFNG is known to induce MNGC formation^102,103^. Our data shows that the primary source of *Ifng* production in the ovary is Type 1 lymphoid cells, which accumulate in the ovary during aging. Additional studies will be required to further elucidate the mechanisms underlying MNGC formation in the ovary and the role they play in ovarian function and aging processes.

The aforementioned accumulation of lymphoid cells and MNGCs almost certainly contribute to pathological changes that occur in the aged ovary. We speculate that they crosstalk with follicular cells to induce immunogenic responses that adversely affect the local microenvironment. For instance, GCs display upregulated IL-17 and IFNG pathway activity with advancing age, although these genes are dominantly expressed in lymphocytes. This supports the notion that lymphoid cells produce pro-inflammatory ligands that alter GC transcriptional networks. We also found that GCs upregulate oxidative stress pathways, which we surmise may occur in response to the pro-inflammatory environment. This is supported by a scRNA-seq analysis of aged primate ovaries which showed declines in oxidoreductase pathway activity in GCs^18^. In addition to immunogenic responses, we also found that GCs and TCs display age-related induction of fibrotic responses as evidenced by increased TGFβ pathway activation. In alignment with prior observations^12,14,15^, we saw increased ovarian collagen deposition by 9 months of age. Since collagen is primarily produced by the stromal fibroblast-like cells^14,104^, we evaluated the expression of collagen genes in this SCL but found no age-related changes. Conversely, the collagenase pathway and MMP2 expression were downregulated in the stroma of aged ovaries, suggesting that collagen accumulation occurs due to declines in degradation. It is also noteworthy that TCs upregulate cell proliferation pathways (e.g. mTOR), which increase with aging^105^, but more importantly have been linked to the activation of primordial follicles^106^.

Although the data presented herein serve as an important tool for hypothesis generation, there are a few limitations that should be noted. Cell preparation for scRNA-seq and flow cytometry requires filtration steps that remove MNGCs and large oocytes from analyses. Future studies employing laser-capture microscopy and single nuclei sequencing would allow for interrogation of these large cellular populations. Even though mice are the most common model organism used for ovarian assessment, there are important aspects of ovarian biology that differ from humans. For example, humans are a mono-ovulating species, whereas mice are poly-ovulating, therefore follicular selection signaling likely differs between these species. In addition, virgin mice were used in this study, which does not account for the impact of pregnancy on ovarian aging dynamics. The examination of additional timepoints following reproductive senescence should also be considered, as it would provide critical insights related to how the ovary changes after the loss of fertility, which may impact systemic aging mechanisms. Finally, estropause in mice does not completely recapitulate the changes observed in human menopause. As such, other models of ovarian aging should be evaluated before definitive conclusions are drawn.

In summary, this report provides insight into changes that occur within the aging murine ovary at single-cell resolution. We report that by 9 months of age, when mice remain fertile and reproductively active, immune cell accumulation is already doubled, with the largest changes being observed in innate lymphocytes such as Type 1 NKTs and Type 17 γδTs. We also demonstrate that GCs and TCs upregulate stress response, immunogenic, and fibrotic signaling pathways with aging. These changes correspond to declines in collagenase expression in the stroma, which we surmise contributes to collagen accumulation with age. These changes were accompanied by accumulation of MNGCs with advancing age, which accounted for the vast majority of lipofuscin positivity in aged ovaries. This observation, coupled with the increase in immune cells that commonly express transcriptional markers of cellular senescence, likely contributes to the previously reported increase in ovarian cellular senescence with advancing age. Collectively, our findings provide insights into the underlying mechanisms that promote chronic ovarian inflammation and fibrosis with advancing age and serve as an important resource for the field.

## METHODS

### Animals and tissue collection and dissociation for scRNA-seq

All animal procedures were approved by the Institutional Animal Care and Use Committee at the Oklahoma Medical Research Foundation (OMRF). C57BL/6J (Strain# 000664) female mice (N=8) were purchased from the Jackson Laboratory (Bar Harbor, ME) and acclimated to the OMRF animal facility before ovary collection. At 3 (n=4) or 9 months of age (n=4), mice were anesthetized with isoflurane and euthanized by exsanguination due to cardiac puncture. Perfusion was performed with 1X PBS, and ovaries were collected and dissected. One ovary from each mouse was dissociated using a Multi Tissue Dissociation Kit 1 (cat# 130110201, Miltenyi Biotec, Bergisch Gladbach, Germany), following manufacturer’s instructions to create a single-cell suspension in D-PBS (cat# 14287080, Gibco).

### scRNA-Seq library construction

scRNA-seq libraries were constructed with the Chromium Single Cell 3ʹ GEM, Library & Gel Bead Kit v3 (cat# 10000075, 10X Genomics), according to the manufacturer’s instructions as briefly described below. Following the creation of ovarian single-cell suspensions, cells were counted on the MACSQuant10 flow cytometer and diluted to 1000 cells per microliter to target the recovery of 5,000 cells per sample during scRNA-seq encapsulation. The diluted cells, master mix, gel beads, and partitioning oil were added to the Chromium Single Cell B Chip (cat# 10000073, 10X Genomics) and loaded into the Chromium controller (cat# 1000204, 10X Genomics) to generate the gel beads-in-emulsion (GEMs) for downstream library preparation. GEMs were then transferred to PCR strip tubes and incubated in a thermocycler to perform reverse transcription in GEMs (GEM-RT). Following GEM-RT, the recovery agent was aspirated, and cDNA was cleaned using the Dynabeads MyOne SILANE reagent included in the scRNA-seq kit. The cDNA was amplified and then cleaned using SPRISelect reagent beads (cat# B23318, Beckman Coulter). The cDNA was quality checked using a High Sensitivity D5000 ScreenTape (cat# 5067-5592, Agilent) run on a TapeStation 2200 (cat# G2964AA, Agilent). An aliquot of 25% of amplified cDNA was carried forward to library preparation. Libraries were quantified by qPCR and quality checked on a High Sensitivity D1000 ScreenTape (cat# 5067-5584, Agilent) on the TapeStation 2220. Libraries were normalized, pooled, and sequenced on the NovaSeq6000 PE150 flow cell. The sequence depth obtained was ∼50,000 reads/ cell.

### scRNA-seq Quality control and data analysis

Fastq files were generated and demultiplexed from raw base call (bcl) files using the cellranger mkfastq pipeline. Alignment, filtering, barcode counting, and UMI counting were conducted using the cellranger count pipeline using the refdata-gex-mm10-2020-A reference transcriptome with default settings. The resultant out files were loaded into R using the “load10X” function of the SoupX package^107^. The SoupX pipeline was then used to estimate and remove ambient RNA contamination before converting to Seurat objects^108^. Samples were then merged to create a single Seurat object and filtered based on the number of features (>200) and percent mitochondrial transcripts (<25%). Genes that were expressed in less than three cells were removed from analysis. Genes that represent ribosomal contamination (*Malat1, Gm42418, Gm26917, Ay036118*)^109,110^ were causing technical background noises and were removed from the analysis to improve sub-clustering. Variable features were identified in Seurat before scaling data and running PCA. The JackStraw method was used to determine the dimensionality of the dataset, and a UMAP analysis was generated in Seurat. Doublets were identified and removed using the DoubletFinder package^111^ with 5% doublets expected. Other parameters (pN=0.25, pK=0.01) were generated using the DoubletFinder sweep statistics. Samples were co-projected on a UMAP which was used to determine that there were no batch effects and that further data integration was not necessary. Seurat was used to find differentially expressed genes by CLU and age and to generate plots presented in the figures (i.e., DimPlot, VlnPlot, DotPlot). Module scores were calculated by the average expression levels of each program on single cell level, subtracted by the aggregated expression of control feature sets. In order to assess cell-to-cell interactions between different ovarian cell types in young and aged murine ovaries, we used CellChat45 (v1.1.3), which is based on the expression of known ligand-receptor pairs^112^.Gene lists were imported into the IPA software Ingenuity Pathway Analysis (IPA) 01.12 (Qiagen Bioinformatics) to assess pathway/biological function enrichment analysis.

### Histology

The remaining ovary from each mouse was collected into 4% PFA, processed, and serially sectioned. To determine ovarian reserve, one of every six serial sections from the whole ovary was H&E stained and follicles from each state were counted as described previously^113^. The number of preantral follicles (primordial, primary, secondary) were multiplied by six to account for sections that were not evaluated and then multiplied by two to account for both ovaries. Due to their large size, antral follicles were multiplied by three to account for sections that were not evaluated and then multiplied by two to account for both ovaries. Picro-Sirius Red staining for collagen deposition and Sudan Black staining for lipofuscin accumulation were performed on one randomly assigned section from the midline of each ovary and analyzed as previously described^11,12^.

### Immunofluorescence

MMP2 protein in the ovarian stroma was evaluated in paraffin-embedded ovarian sections as previously described^114^. The slides were incubated with a primary rabbit anti-MMP2 antibody (cat# 87809, BioLegend, San Diego, CA; 1:100) overnight, followed by a secondary goat anti-rabbit IgG Alexa Fluor 488 antibody (cat# 2338052, Jackson ImmunoResearch Laboratories; 1:500) for 1h and DAPI for 5 min. Images were captured on a fluorescent microscope (Zeiss, Oberkochen, Germany) in the green (MMP2) and blue (DAPI) channels. Images were also taken in the red channel to identify and avoid autofluorescent regions in the ovary, which are common in aged ovaries^43^. The mean fluorescent intensity from three MMP2-stained stromal regions from each mouse were quantified by Image J and averaged. Nonspecific background fluorescence was calculated in a similar manner using secondary-only stained serial sections from the same mouse. The nonspecific signal was subtracted from each sample.

### Flow Cytometry

High parameter spectral flow cytometry was performed to confirm age-related changes in the percentage and numbers of immune cells. C57BL/6J (Strain# 000664) female mice (N=30) were purchased from the Jackson Laboratory. To obtain enough cells for the gating strategy proposed, six ovaries from three mice were pooled (n=5/age). Prior to euthanasia, vaginal cytology was performed to determine estrous cycle stage as previously described^92^. The mice were anesthetized with isoflurane and intravenously (i.v.) injected with 2 μg of FITC-anti-CD45 mAb (cat# 35-0454, Cytek, San Diego, CA) to label intravascular cells and distinguish them from tissue-resident extravascular leukocytes^115^. Five minutes after labeling, mice were euthanized and ovaries were collected. Ovaries were pooled according to estrous cycle stage. Ovarian cells were isolated by enzymatic digestion in 3 mL of Dulbecco’s Modified Eagle’s Medium (DMEM) (cat# D6429, Sigma) containing 4 mg collagenase (cat# C5138-100MG, Sigma-Aldrich, St. Louis, MO). Samples were incubated at 37°C for 40 minutes and gently pipetted 30 times every 10 minutes to encourage tissue dissociation^93^. Following dissociation, cells were passed through a 70 µm filter (cat# 130-098-462, Miltenyi Biotec) and washed with an additional 7 mL of DMEM. Cells were labeled with Zombie NIR (cat# 423106, Biolegend, San Diego, CA, USA) according to the manufacturer’s instructions, then washed in FACS Buffer (PBS + 5% Newborn Calf Serum), incubated with a Fc blocking reagent (anti-mouse CD16/32; cat# 70-0161, Cytek), washed and stained with a surface staining fluorochrome-labeled mAb cocktail for 30 min at 4° C, in the presence of Brilliant Stain Buffer Plus (10 μl/sample) (cat# 563795, Becton Dickinson, Mountain View, Ca, USA) and CellBlox Blocking Buffer (5 μl/sample) (cat# C001T02F01, Thermo Fisher Scientific). Cells were washed again in FACS buffer and intracellularly stained to detect transcription factor expression using the True-Nuclear Transcription Factor Buffer Set (cat# 424401, Biolegend) according to the manufacturer’s instructions. At the end of the procedure, stained cells were fixed in 2% paraformaldehyde for 5 min. Stained cells were acquired using a 5-laser Cytek Aurora flow cytometer, and analyzed using FlowJo 10.9 (Becton Dickinson). The gating strategy can be seen in Suppl. Fig. 4.

### Statistics

Differentially expressed genes (DEGs) between CLU and by age were called in the Seurat package, using the FindMarkers command with default parameters. DEG lists were imported into IPA software and filtered on FDR<0.1 and logFC>0.25 for pathway analyses. Pathways with p<0.05 were considered statistically significant and the activation z-scores were reported by heatmap or bar charts in the figures. More traditional statistical analyses were done using GraphPad Prism software. Strip plots are presented with individual points shown and means ± SEM indicated. For comparisons of means between the two ages, Student’s t-tests were used, applying one- and two-tailed tests where appropriate^116^. Corrections for multiple comparisons were made, where appropriate, using the Benjamini, Krieger, and Yekutieli correction for multiple comparisons. Significant differences were defined at P<0.05 or FDR<0.05 (for multiple comparisons).

## ACKNOWLEDGEMENTS

This work was supported by the National Institutes of Health (R01 AG069742 to M.B.S. and P30 AG050911 to W.M.F.), Global Consortium for Reproductive Longevity and Equality (GCRLE-4501 to M.B.S and GCRLE-0523 to S.R.O.) and Presbyterian Health Foundation (Pilot Research Funding to M.B.S.). The authors thank the staff at the Clinical Genomics Center, Flow Cytometry Core (Cytek Aurora, Grant number: 1S10OD028479-01), and Imaging Core Facility at the Oklahoma Medical Research Foundation for assistance with sequencing and histological procedures.

## AUTHOR CONTRIBUTIONS

J.V.V.I., S.R.O., and M.B.S. conceived the project and designed the experiments. J.V.V.I. and S.R.O. performed the experiments and data analyses with contributions from C.R.H., S.K., S.A.M., J.D.H., H.N.C.C., A.S., S.K., J.A.I. and W.M.F. J.V.V.I., S.R.O., and C.R.H. created figures and performed statistical analyses with contributions from W.M.F. and M.B.S. J.V.V.I., S.R.O., and M.B.S. wrote the manuscript and all authors edited and approved the final draft.

## DATA AVAILABILITY STATEMENT

The datasets generated through this work are available upon reasonable request from the corresponding author.

## COMPLETING INTERESTS

The authors declare no conflicts or competing interests.

---

**Suppl. Fig. 1.**
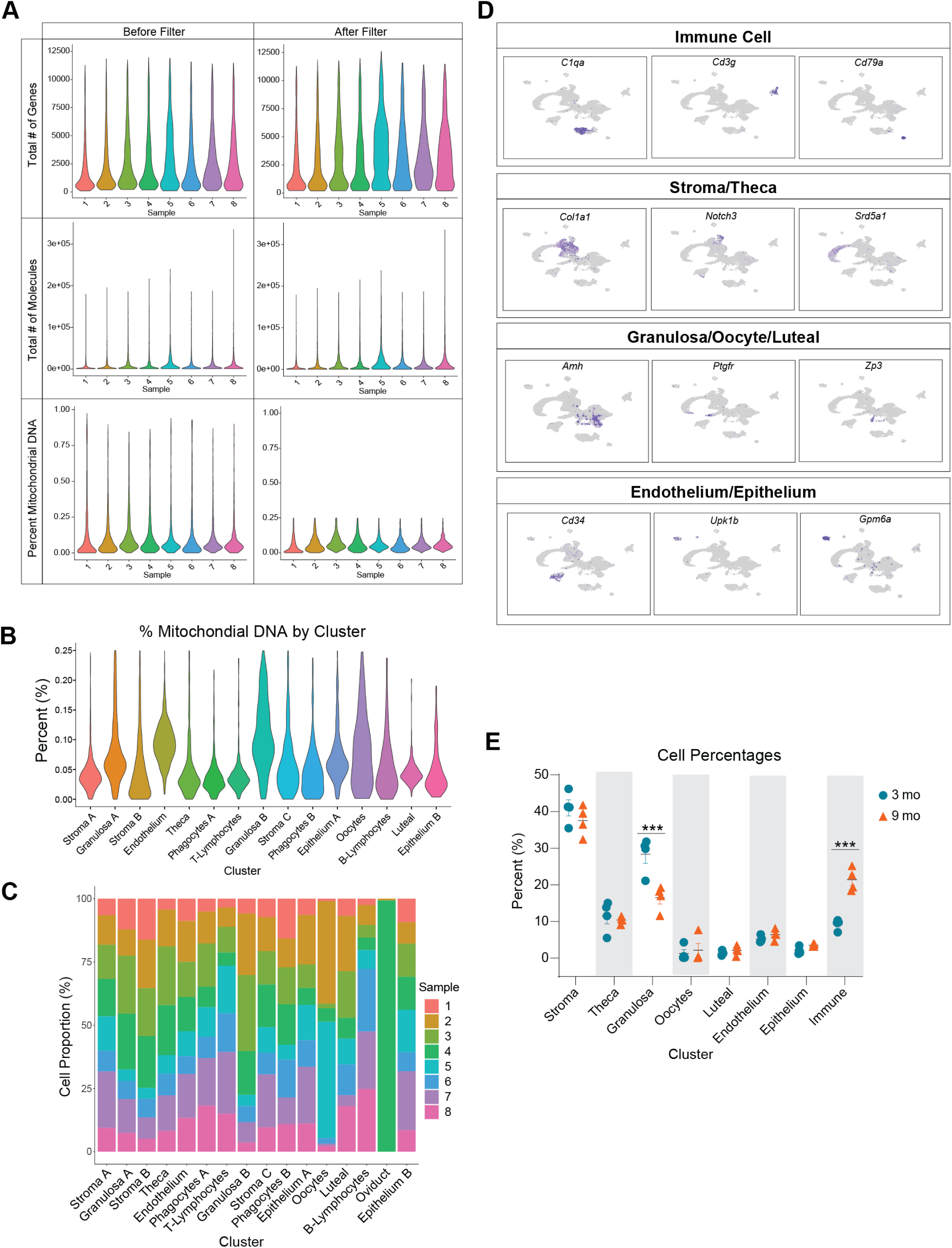
(A) Violin plots of the total number of genes and molecules and mitochondrial percentages within each sample before and after quality filtering. (B) Percent of mitochondrial RNA by CLU. (C) Percentage of cells in each CLU by sample, showing oviduct cells contamination in one sample which was removed from further analyses. (D) Feature plots of specific marker genes of cell types used to identify the CLUs. (E) Cell percentages in each broad cell type by sample. scRNA-seq was performed in n=4 ovaries/age. Data are presented as mean ± SEM. *, **, *** represent statistical difference (FDR<0.05, 0.01 and 0.005, respectively) by multiple two-tailed t-test with Benjamini, Krieger, and Yekutieli correction for multiple comparisons.

**Suppl. Fig. 2.**
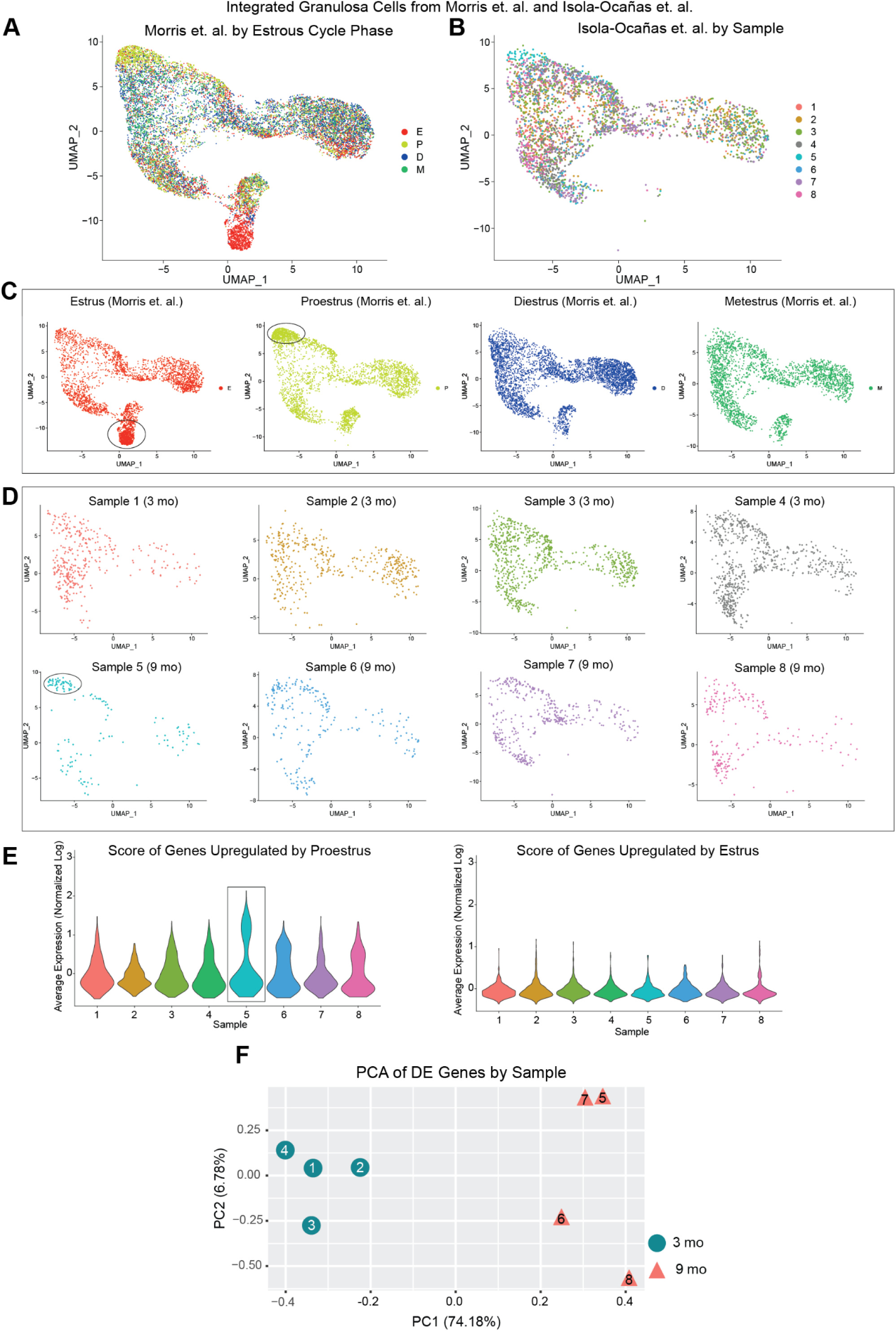
*In silico* estrous cycle staging by comparison to Morris et al., 2022. GCs from the Morris et al. study were co-clustered with GC from the present study to infer estrous cycle stage. (A) UMAP plot featuring GC across different estrous cycle stages from Morris et al. (B) UMAP plot featuring GC from all samples of the present study integrated with Morris et al. dataset. (C) UMAP plots from Morris et al. featuring each phase of estrous cycle, noting distinct CLUs only observed during the estrus and proestrus phases. (D) UMAP plot featuring GC from each samples of the present study, suggesting Sample 5 is in proestrus and the remaining samples are in either metestrus or diestrus. (E) Module score of genes reported by Morris et al. as upregulated in GC in both estrus and proestrus in each of our samples, suggesting Sample 5 is in proestrus and the remaining samples are in either metestrus or diestrus. (F) PCA of gene expression by sample, showing clear separation of samples by age. scRNA-seq was performed in n=4 ovaries/age.

**Suppl. Fig. 3.**
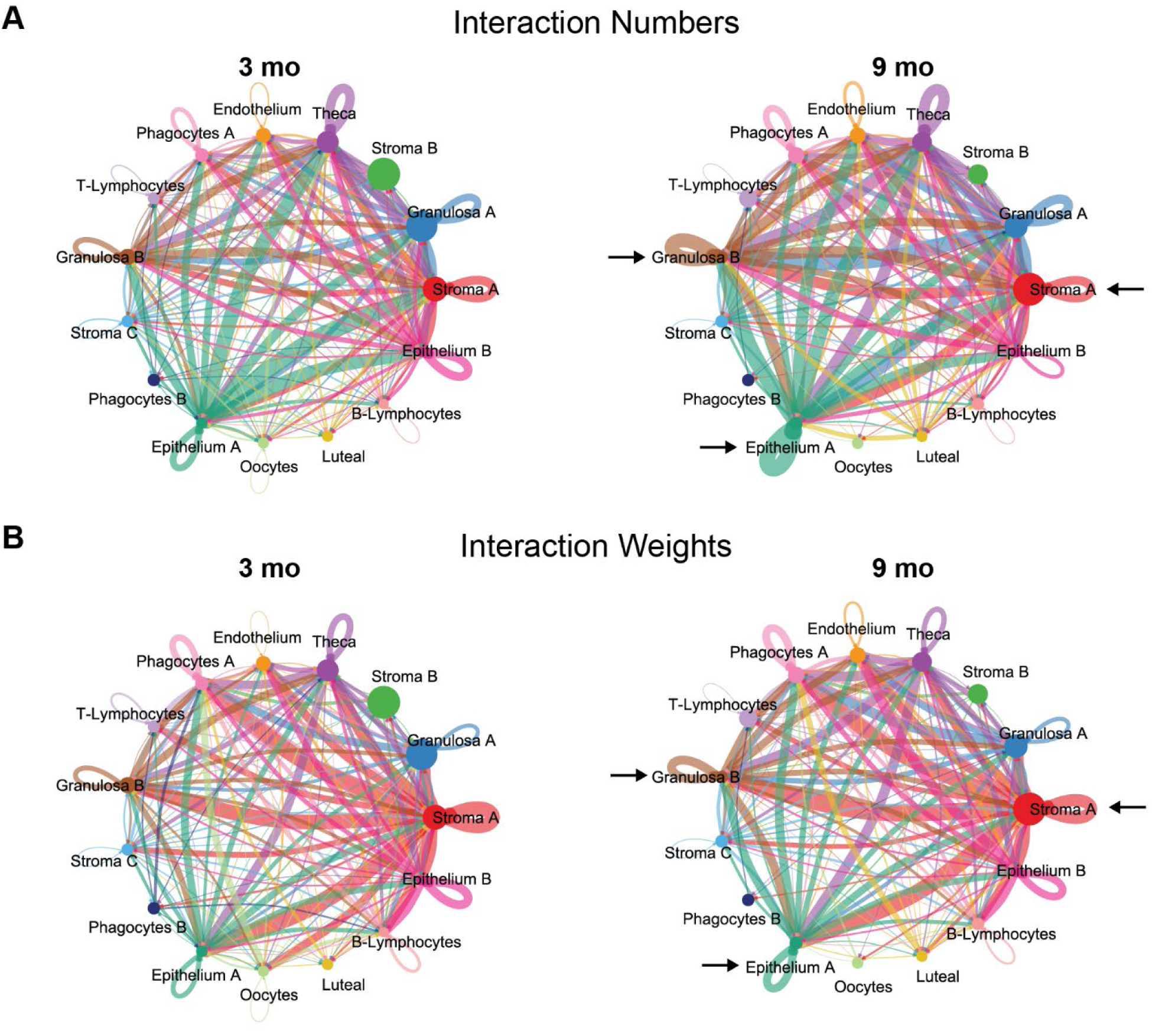
Cellular communication networks in the aging murine ovary. (A) Number of cellular interactions in the 3-mo and 9-mo ovarian using the CellChat signaling package. (B) Interaction weights in the 3-mo and 9-mo ovarian using the CellChat signaling package. scRNA-seq was performed in n=4 ovaries/age.

**Suppl. Fig. 4.**
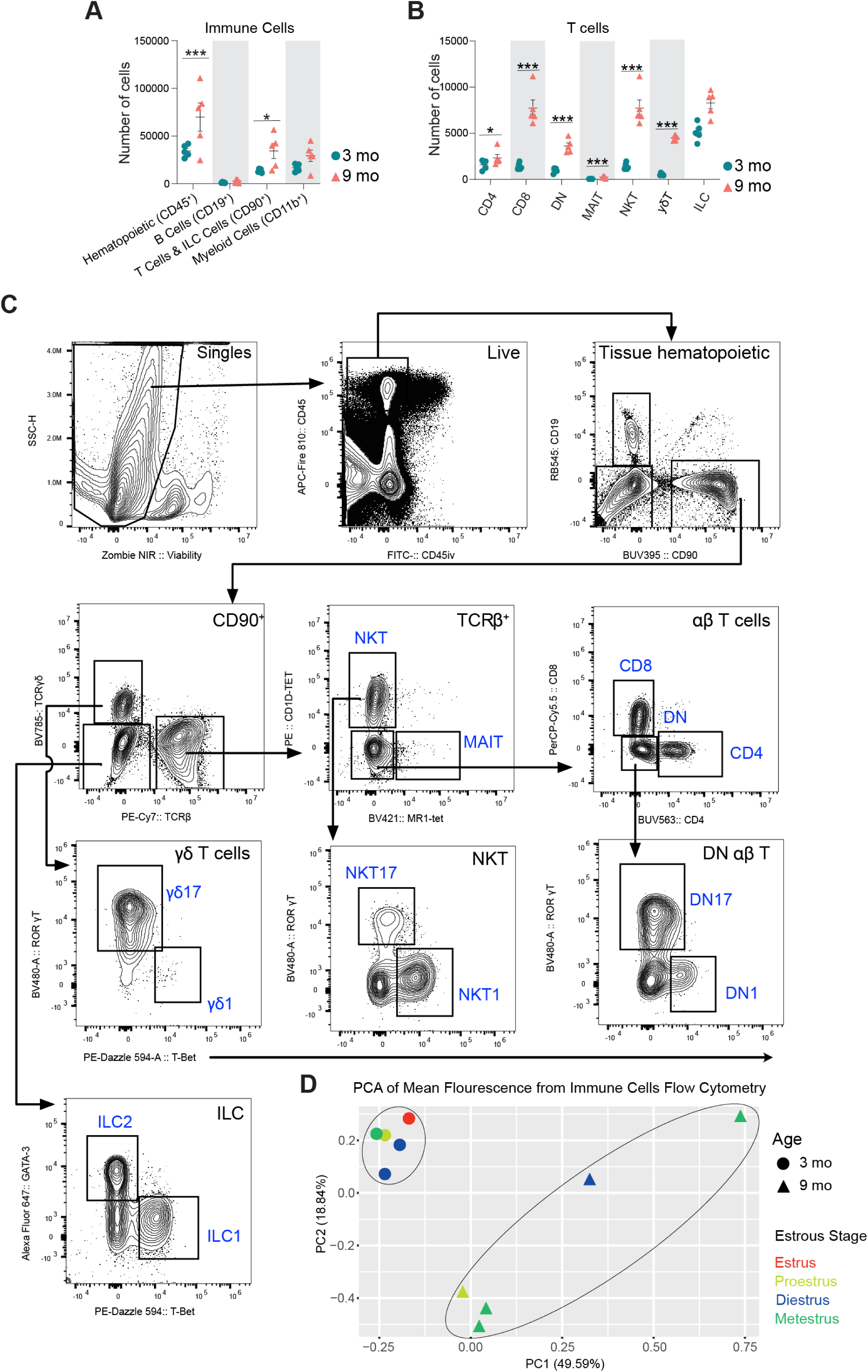
(A-B) Absolute numbers of ovarian immune cells by flow cytometry after gating out intravascular hematopoietic cells. (C) Gating strategy for lymphocyte flow cytometry. (D) PCA of mean fluorescence by channel in immune cells flow cytometry, showing separation by age and not estrous cycle stage. For flow cytometry, 6 ovaries from mice in the same phase of estrous cycle were pooled together to comprise each sample (n=5/age). Data are presented as mean ± SEM. *, **, *** represent statistical difference (FDR<0.05, 0.01 and 0.005, respectively) by multiple two-tailed t-test with Benjamini, Krieger, and Yekutieli correction for multiple comparisons.

**Suppl. Fig. 5.**
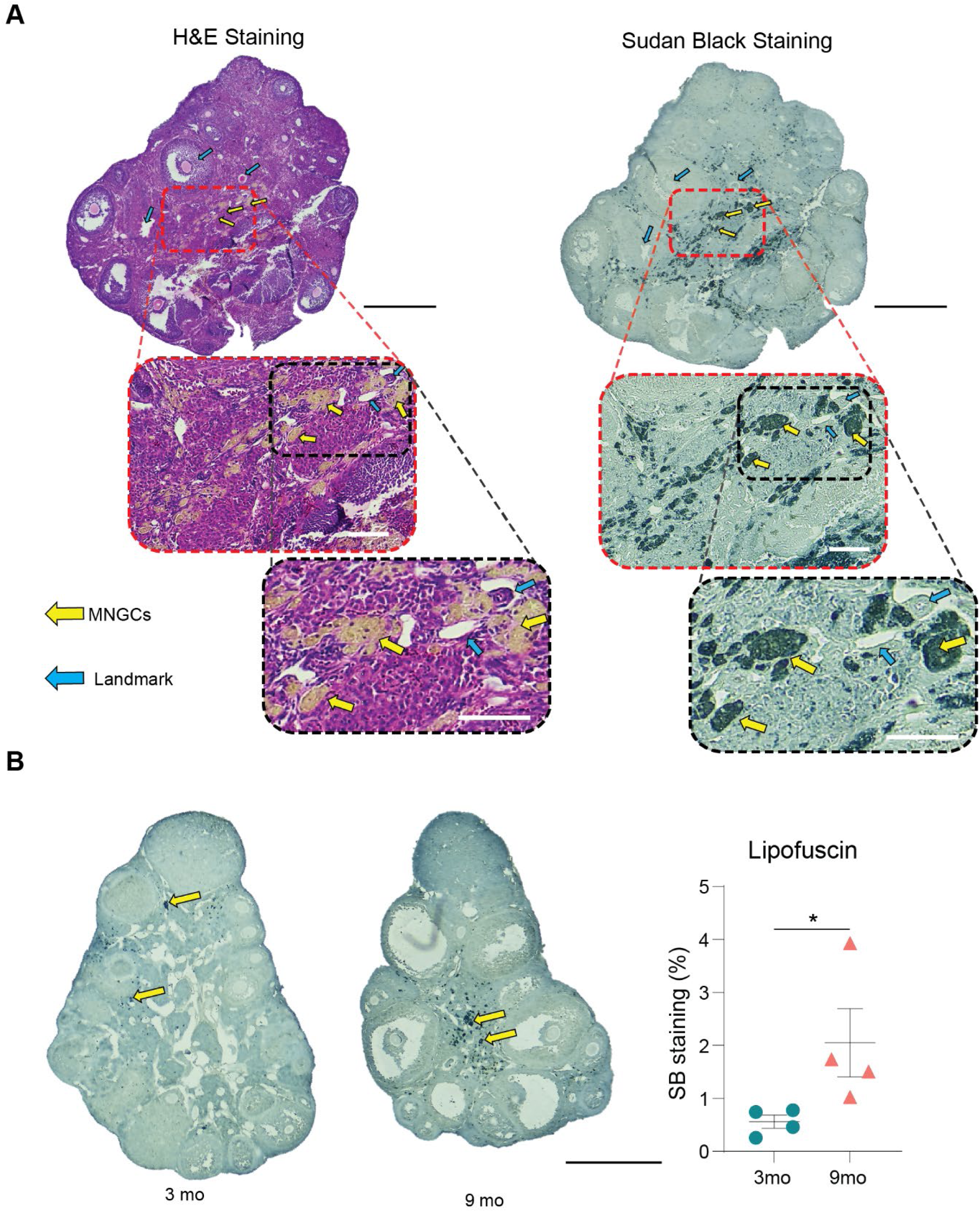
Assessment of lipofuscin accumulation in the aged ovary. (A) H&E and Sudan black staining of 9-month-old ovarian sections showing MNGC accumulation compared to 3-month-old ovaries. (B) Sudan black staining of lipofuscin in 3- and 9-month-old ovaries. Data are presented as mean ± SEM. * represents statistical difference (p<0.05) by one-tailed t-test. scRNA-seq was performed in n=4 ovaries/age. Black scale bars represent 500µm and white scale bars represent 100µm.

**Suppl. Fig. 6.**
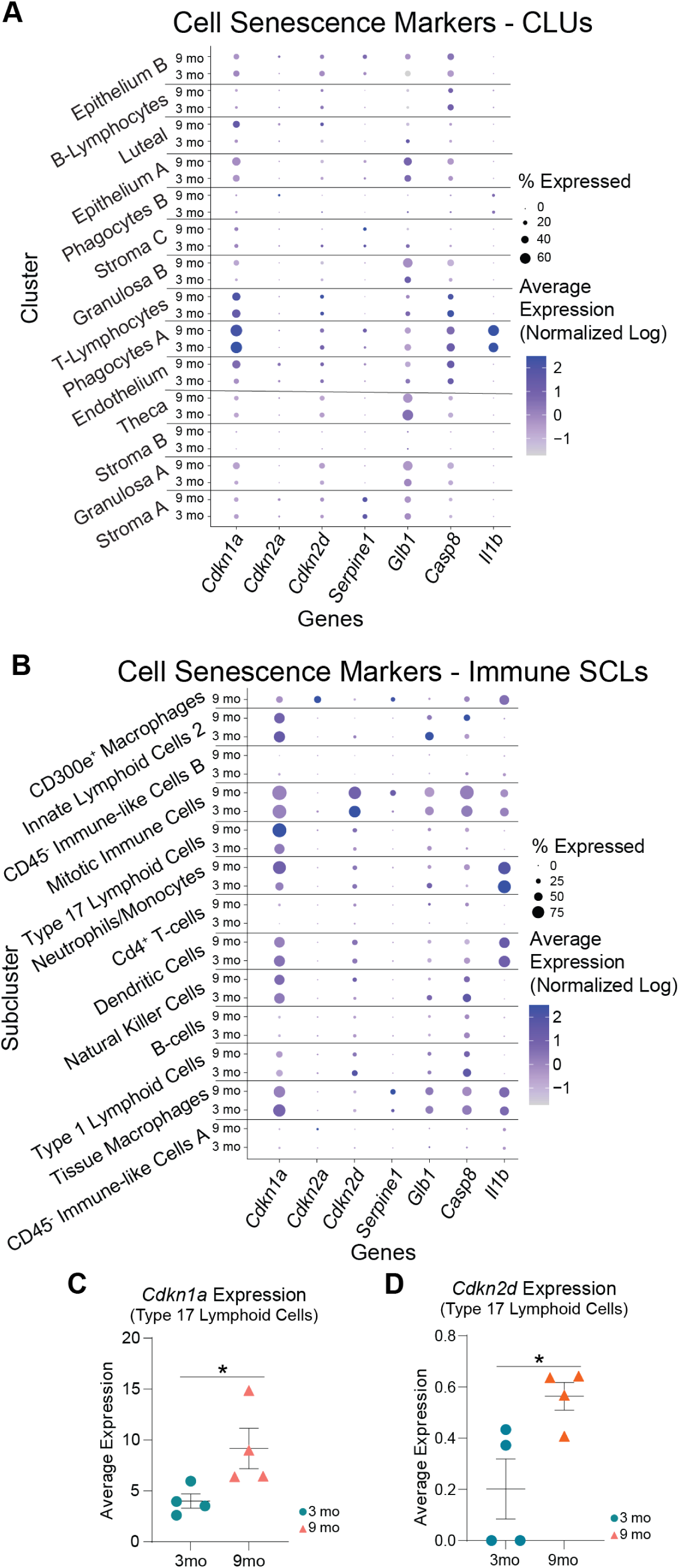
Expression of cellular senescence markers is unchanged with age. (A) Dot plot of expression of cellular senescence markers in initial CLUs. (B) Dot plot of expression of cell senescence markers in immune SCLs. (C) *Cdnk1a* expression in Type 17 lymphoid cells. scRNA-seq was performed in n=4 ovaries/age. Data are presented as mean ± SEM. *, **, *** represent statistical difference (p<0.05, 0.01 and 0.005, respectively) by one-tailed t-test.

**Suppl. Fig. 7:**
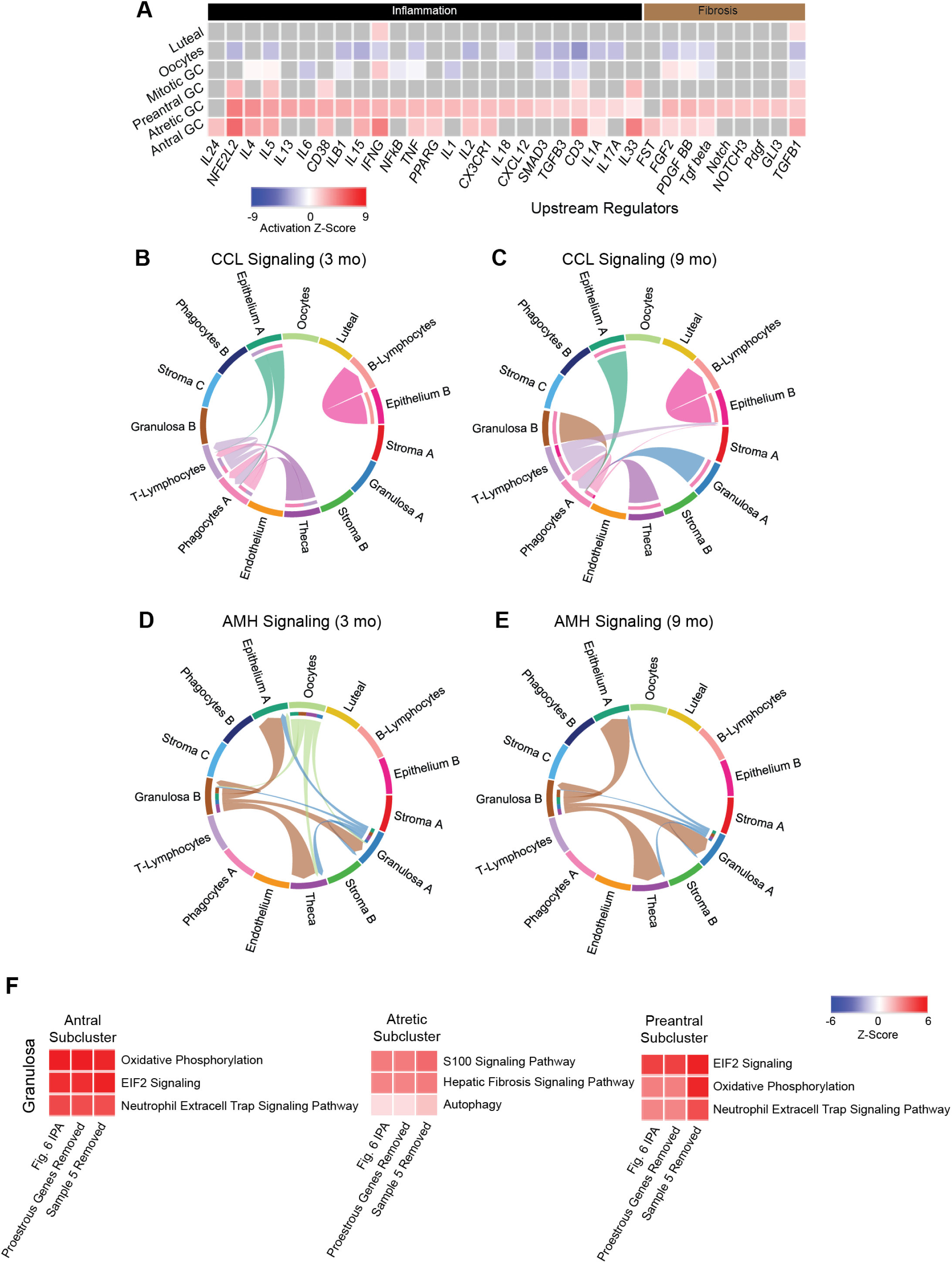
Cell signaling in granulosa cells, oocytes and luteal cells. (A) Z-scores indicating activation or inhibition of upstream regulators with aging by IPA analysis in granulosa/oocytes/luteal SCLs. (B-C) CellChat chord diagrams of the CCL signaling pathway interactions in the 3 and 9 month-old ovarian CLUs. (D-E) CellChat chord diagrams of the AMH signaling pathway interactions in the 3 and 9 month-old ovarian CLUs. (F) Heatmaps of three of the most changed IPA pathways from Fig. 6 in granulosa SCLs with and without the removal of proestrus upregulated genes or sample 5, showing little to no changes. scRNA-seq was performed in n=4 ovaries/age.

## REFERENCES

1 Johnson, J. A. & Tough, S. No-271-Delayed Child-Bearing. J Obstet Gynaecol Can 39, e500–e515, doi:10.1016/j.jogc.2017.09.007 (2017).

2 Broekmans, F. J., Soules, M. R. & Fauser, B. C. Ovarian aging: mechanisms and clinical consequences. Endocr Rev 30, 465–493, doi:10.1210/er.2009-0006 (2009).

3 Levine, M. E. et al. Menopause accelerates biological aging. Proc Natl Acad Sci U S A 113, 9327–9332, doi:10.1073/pnas.1604558113 (2016).

4 Wellons, M., Ouyang, P., Schreiner, P. J., Herrington, D. M. & Vaidya, D. Early menopause predicts future coronary heart disease and stroke: the Multi-Ethnic Study of Atherosclerosis. Menopause 19, 1081–1087, doi:10.1097/gme.0b013e3182517bd0 (2012).

5 Tchernof, A., Calles-Escandon, J., Sites, C. K. & Poehlman, E. T. Menopause, central body fatness, and insulin resistance: effects of hormone-replacement therapy. Coron Artery Dis 9, 503–511, doi:10.1097/00019501-199809080-00006 (1998).

6 Muka, T. et al. Association of Age at Onset of Menopause and Time Since Onset of Menopause With Cardiovascular Outcomes, Intermediate Vascular Traits, and All-Cause Mortality: A Systematic Review and Meta-analysis. JAMA Cardiol 1, 767–776, doi:10.1001/jamacardio.2016.2415 (2016).

7 Ossewaarde, M. E. et al. Age at menopause, cause-specific mortality and total life expectancy. Epidemiology 16, 556–562, doi:10.1097/01.ede.0000165392.35273.d4 (2005).

8 May-Panloup, P. et al. Ovarian ageing: the role of mitochondria in oocytes and follicles. Hum Reprod Update 22, 725–743, doi:10.1093/humupd/dmw028 (2016).

9 Lim, J. & Luderer, U. Oxidative damage increases and antioxidant gene expression decreases with aging in the mouse ovary. Biol Reprod 84, 775–782, doi:10.1095/biolreprod.110.088583 (2011).

10 Yang, L. et al. The Role of Oxidative Stress and Natural Antioxidants in Ovarian Aging. Front Pharmacol 11, 617843, doi:10.3389/fphar.2020.617843 (2020).

11 Ansere, V. A. et al. Cellular hallmarks of aging emerge in the ovary prior to primordial follicle depletion. Mech Ageing Dev 194, 111425, doi:10.1016/j.mad.2020.111425 (2021).

12 Briley, S. M. et al. Reproductive age-associated fibrosis in the stroma of the mammalian ovary. Reproduction 152, 245–260, doi:10.1530/REP-16-0129 (2016).

13 Lliberos, C. et al. Evaluation of inflammation and follicle depletion during ovarian ageing in mice. Sci Rep 11, 278, doi:10.1038/s41598-020-79488-4 (2021).

14 Amargant, F. et al. Ovarian stiffness increases with age in the mammalian ovary and depends on collagen and hyaluronan matrices. Aging Cell 19, e13259, doi:10.1111/acel.13259 (2020).

15 Umehara, T. et al. Female reproductive life span is extended by targeted removal of fibrotic collagen from the mouse ovary. Sci Adv 8, eabn4564, doi:10.1126/sciadv.abn4564 (2022).

16 Mara, J. N. et al. Ovulation and ovarian wound healing are impaired with advanced reproductive age. Aging (Albany NY) 12, 9686–9713, doi:10.18632/aging.103237 (2020).

17 Foley, K. G., Pritchard, M. T. & Duncan, F. E. Macrophage-derived multinucleated giant cells: hallmarks of the aging ovary. Reproduction 161, V5–V9, doi:10.1530/REP-20-0489 (2021).

18 Wang, S. et al. Single-Cell Transcriptomic Atlas of Primate Ovarian Aging. Cell 180, 585–600 e519, doi:10.1016/j.cell.2020.01.009 (2020).

19 Wang, S. et al. Spatiotemporal analysis of human ovarian aging at single-cell resolution. Research Square, doi:10.21203/rs.3.rs-1624864/v1 (2022).

20 Lu, H. et al. Current Animal Model Systems for Ovarian Aging Research. Aging Dis 13, 1183–1195, doi:10.14336/AD.2021.1209 (2022).

21 Russ, J. E., Haywood, M. E., Lane, S. L., Schoolcraft, W. B. & Katz-Jaffe, M. G. Spatially resolved transcriptomic profiling of ovarian aging in mice. iScience 25, 104819, doi:10.1016/j.isci.2022.104819 (2022).

22 Ankel-Simons, F. & Cummins, J. M. Misconceptions about mitochondria and mammalian fertilization: implications for theories on human evolution. Proc Natl Acad Sci U S A 93, 13859–13863, doi:10.1073/pnas.93.24.13859 (1996).

23 Zhang, D., Keilty, D., Zhang, Z. F. & Chian, R. C. Mitochondria in oocyte aging: current understanding. Facts Views Vis Obgyn 9, 29–38 (2017).

24 Zhang, G. et al. Expression of Mitochondria-Associated Genes (PPARGC1A, NRF-1, BCL-2 and BAX) in Follicular Development and Atresia of Goat Ovaries. Reprod Domest Anim 50, 465–473, doi:10.1111/rda.12514 (2015).

25 Muhl, L. et al. Single-cell analysis uncovers fibroblast heterogeneity and criteria for fibroblast and mural cell identification and discrimination. Nat Commun 11, 3953, doi:10.1038/s41467-020-17740-1 (2020).

26 Morris, M. E. et al. A single-cell atlas of the cycling murine ovary. Elife 11, doi:10.7554/eLife.77239 (2022).

27 Baek, S. H. et al. Single Cell Transcriptomic Analysis Reveals Organ Specific Pericyte Markers and Identities. Front Cardiovasc Med 9, 876591, doi:10.3389/fcvm.2022.876591 (2022).

28 Wagner, M. et al. Single-cell analysis of human ovarian cortex identifies distinct cell populations but no oogonial stem cells. Nat Commun 11, 1147, doi:10.1038/s41467-020-14936-3 (2020).

29 Marti, N. et al. Genes and proteins of the alternative steroid backdoor pathway for dihydrotestosterone synthesis are expressed in the human ovary and seem enhanced in the polycystic ovary syndrome. Mol Cell Endocrinol 441, 116–123, doi:10.1016/j.mce.2016.07.029 (2017).

30 Sontheimer, R. D., Racila, E. & Racila, D. M. C1q: its functions within the innate and adaptive immune responses and its role in lupus autoimmunity. J Invest Dermatol 125, 14–23, doi:10.1111/j.0022-202X.2005.23673.x (2005).

31 Lin, G., Finger, E. & Gutierrez-Ramos, J. C. Expression of CD34 in endothelial cells, hematopoietic progenitors and nervous cells in fetal and adult mouse tissues. Eur J Immunol 25, 1508–1516, doi:10.1002/eji.1830250606 (1995).

32 Garcillan, B. et al. CD3G or CD3D Knockdown in Mature, but Not Immature, T Lymphocytes Similarly Cripples the Human TCRalphabeta Complex. Front Cell Dev Biol 9, 608490, doi:10.3389/fcell.2021.608490 (2021).

33 Carpenter, A. R. et al. Uroplakin 1b is critical in urinary tract development and urothelial differentiation and homeostasis. Kidney Int 89, 612–624, doi:10.1016/j.kint.2015.11.017 (2016).

34 Berisha, B., Rodler, D., Schams, D., Sinowatz, F. & Pfaffl, M. W. Prostaglandins in Superovulation Induced Bovine Follicles During the Preovulatory Period and Early Corpus Luteum. Front Endocrinol (Lausanne) 10, 467, doi:10.3389/fendo.2019.00467 (2019).

35 Chu, P. G. & Arber, D. A. CD79: a review. Appl Immunohistochem Mol Morphol 9, 97–106, doi:10.1097/00129039-200106000-00001 (2001).

36 Hu, C. et al. CellMarker 2.0: an updated database of manually curated cell markers in human/mouse and web tools based on scRNA-seq data. Nucleic Acids Res 51, D870–D876, doi:10.1093/nar/gkac947 (2023).

37 Heng, T. S., Painter, M. W. & Immunological Genome Project, C. The Immunological Genome Project: networks of gene expression in immune cells. Nat Immunol 9, 1091–1094, doi:10.1038/ni1008-1091 (2008).

38 Szabo, S. J. et al. A novel transcription factor, T-bet, directs Th1 lineage commitment. Cell 100, 655–669, doi:10.1016/s0092-8674(00)80702-3 (2000).

39 Ivanov, II et al. The orphan nuclear receptor RORgammat directs the differentiation program of proinflammatory IL-17+ T helper cells. Cell 126, 1121–1133, doi:10.1016/j.cell.2006.07.035 (2006).

40 Savage, A. K. et al. The transcription factor PLZF directs the effector program of the NKT cell lineage. Immunity 29, 391–403, doi:10.1016/j.immuni.2008.07.011 (2008).

41 Kovalovsky, D. et al. The BTB-zinc finger transcriptional regulator PLZF controls the development of invariant natural killer T cell effector functions. Nat Immunol 9, 1055–1064, doi:10.1038/ni.1641 (2008).

42 Coletta, S. et al. The immune receptor CD300e negatively regulates T cell activation by impairing the STAT1-dependent antigen presentation. Sci Rep 10, 16501, doi:10.1038/s41598-020-73552-9 (2020).

43 Asano, Y. Age-related accumulation of non-heme ferric and ferrous iron in mouse ovarian stroma visualized by sensitive non-heme iron histochemistry. J Histochem Cytochem 60, 229–242, doi:10.1369/0022155411431734 (2012).

44 Urzua, U., Chacon, C., Espinoza, R., Martinez, S. & Hernandez, N. Parity-Dependent Hemosiderin and Lipofuscin Accumulation in the Reproductively Aged Mouse Ovary. Anal Cell Pathol (Amst) 2018, 1289103, doi:10.1155/2018/1289103 (2018).

45 Evangelou, K. & Gorgoulis, V. G. Sudan Black B, The Specific Histochemical Stain for Lipofuscin: A Novel Method to Detect Senescent Cells. Methods Mol Biol 1534, 111–119, doi:10.1007/978-1-4939-6670-7_10 (2017).

46 Ruan, J. et al. Novel Myh11 Dual Reporter Mouse Model Provides Definitive Labeling and Identification of Smooth Muscle Cells-Brief Report. Arterioscler Thromb Vasc Biol 41, 815–821, doi:10.1161/ATVBAHA.120.315107 (2021).

47 Deaton, R. A. et al. A New Autosomal Myh11-CreER(T2) Smooth Muscle Cell Lineage Tracing and Gene Knockout Mouse Model-Brief Report. Arterioscler Thromb Vasc Biol 43, 203–211, doi:10.1161/ATVBAHA.122.318160 (2023).

48 Rosas-Canyelles, E., Dai, T., Li, S. & Herr, A. E. Mouse-to-mouse variation in maturation heterogeneity of smooth muscle cells. Lab Chip 18, 1875–1883, doi:10.1039/c8lc00216a (2018).

49 Martinez, F. O., Helming, L. & Gordon, S. Alternative activation of macrophages: an immunologic functional perspective. Annu Rev Immunol 27, 451–483, doi:10.1146/annurev.immunol.021908.132532 (2009).

50 Steiger, S. et al. Immunomodulatory Molecule IRAK-M Balances Macrophage Polarization and Determines Macrophage Responses during Renal Fibrosis. J Immunol 199, 1440–1452, doi:10.4049/jimmunol.1601982 (2017).

51 Li, Y. L., Sato, M., Kojima, N., Miura, M. & Senoo, H. Regulatory role of extracellular matrix components in expression of matrix metalloproteinases in cultured hepatic stellate cells. Cell Struct Funct 24, 255–261, doi:10.1247/csf.24.255 (1999).

52 Nelson, J. F., Felicio, L. S., Randall, P. K., Sims, C. & Finch, C. E. A longitudinal study of estrous cyclicity in aging C57BL/6J mice: I. Cycle frequency, length and vaginal cytology. Biol Reprod 27, 327–339, doi:10.1095/biolreprod27.2.327 (1982).

53 Knight, P. G. & Glister, C. TGF-beta superfamily members and ovarian follicle development. Reproduction 132, 191–206, doi:10.1530/rep.1.01074 (2006).

54 McCloskey, C. W. et al. Metformin Abrogates Age-Associated Ovarian Fibrosis. Clin Cancer Res 26, 632–642, doi:10.1158/1078-0432.CCR-19-0603 (2020).

55 Li, M. O., Wan, Y. Y., Sanjabi, S., Robertson, A. K. & Flavell, R. A. Transforming growth factor-beta regulation of immune responses. Annu Rev Immunol 24, 99–146, doi:10.1146/annurev.immunol.24.021605.090737 (2006).

56 Dompe, C. et al. Human Granulosa Cells-Stemness Properties, Molecular Cross-Talk and Follicular Angiogenesis. Cells 10, doi:10.3390/cells10061396 (2021).

57 Meinsohn, M. C. et al. Single-cell sequencing reveals suppressive transcriptional programs regulated by MIS/AMH in neonatal ovaries. Proc Natl Acad Sci U S A 118, doi:10.1073/pnas.2100920118 (2021).

58 Fan, X. et al. Single-cell reconstruction of follicular remodeling in the human adult ovary. Nat Commun 10, 3164, doi:10.1038/s41467-019-11036-9 (2019).

59 Chen, A. Q., Wang, Z. G., Xu, Z. R., Yu, S. D. & Yang, Z. G. Analysis of gene expression in granulosa cells of ovine antral growing follicles using suppressive subtractive hybridization. Anim Reprod Sci 115, 39–48, doi:10.1016/j.anireprosci.2008.10.022 (2009).

60 Wigglesworth, K., Lee, K. B., Emori, C., Sugiura, K. & Eppig, J. J. Transcriptomic diversification of developing cumulus and mural granulosa cells in mouse ovarian follicles. Biol Reprod 92, 23, doi:10.1095/biolreprod.114.121756 (2015).

61 Lee, J. H. & Berger, J. M. Cell Cycle-Dependent Control and Roles of DNA Topoisomerase II. Genes (Basel) 10, doi:10.3390/genes10110859 (2019).

62 Blanchard, J. M. Cyclin A2 transcriptional regulation: modulation of cell cycle control at the G1/S transition by peripheral cues. Biochem Pharmacol 60, 1179–1184, doi:10.1016/s0006-2952(00)00384-1 (2000).

63 Piersanti, R. L., Santos, J. E. P., Sheldon, I. M. & Bromfield, J. J. Lipopolysaccharide and tumor necrosis factor-alpha alter gene expression of oocytes and cumulus cells during bovine in vitro maturation. Mol Reprod Dev 86, 1909–1920, doi:10.1002/mrd.23288 (2019).

64 Stocco, C., Telleria, C. & Gibori, G. The molecular control of corpus luteum formation, function, and regression. Endocr Rev 28, 117–149, doi:10.1210/er.2006-0022 (2007).

65 Salmon, N. A., Handyside, A. H. & Joyce, I. M. Oocyte regulation of anti-Mullerian hormone expression in granulosa cells during ovarian follicle development in mice. Dev Biol 266, 201–208, doi:10.1016/j.ydbio.2003.10.009 (2004).

66 Shrikhande, L., Shrikhande, B. & Shrikhande, A. AMH and Its Clinical Implications. J Obstet Gynaecol India 70, 337–341, doi:10.1007/s13224-020-01362-0 (2020).

67 Brown, H. M. & Russell, D. L. Blood and lymphatic vasculature in the ovary: development, function and disease. Hum Reprod Update 20, 29–39, doi:10.1093/humupd/dmt049 (2014).

68 Brown, H. M., Robker, R. L. & Russell, D. L. Development and hormonal regulation of the ovarian lymphatic vasculature. Endocrinology 151, 5446–5455, doi:10.1210/en.2010-0629 (2010).

69 Hartanti, M. D. et al. Formation of the Bovine Ovarian Surface Epithelium during Fetal Development. J Histochem Cytochem 68, 113–126, doi:10.1369/0022155419896797 (2020).

70 Hummitzsch, K. et al. A new model of development of the mammalian ovary and follicles. PLoS One 8, e55578, doi:10.1371/journal.pone.0055578 (2013).

71 Schulz, A. et al. The Soluble Fms-like Tyrosine Kinase-1 Contributes to Structural and Functional Changes in Endothelial Cells in Chronic Kidney Disease. Int J Mol Sci 23, doi:10.3390/ijms232416059 (2022).

72 Galvagni, F. et al. Dissecting the CD93-Multimerin 2 interaction involved in cell adhesion and migration of the activated endothelium. Matrix Biol 64, 112–127, doi:10.1016/j.matbio.2017.08.003 (2017).

73 Fujimoto, N. et al. Single-cell mapping reveals new markers and functions of lymphatic endothelial cells in lymph nodes. PLoS Biol 18, e3000704, doi:10.1371/journal.pbio.3000704 (2020).

74 Bloom, S. I., Islam, M. T., Lesniewski, L. A. & Donato, A. J. Mechanisms and consequences of endothelial cell senescence. Nat Rev Cardiol 20, 38–51, doi:10.1038/s41569-022-00739-0 (2023).

75 Wang, J. J. et al. Single-cell transcriptome landscape of ovarian cells during primordial follicle assembly in mice. PLoS Biol 18, e3001025, doi:10.1371/journal.pbio.3001025 (2020).

76 Pei, J. et al. Single-Cell Transcriptomics Analysis Reveals a Cell Atlas and Cell Communication in Yak Ovary. Int J Mol Sci 24, doi:10.3390/ijms24031839 (2023).

77 Gilardi, K. V., Shideler, S. E., Valverde, C. R., Roberts, J. A. & Lasley, B. L. Characterization of the onset of menopause in the rhesus macaque. Biol Reprod 57, 335–340, doi:10.1095/biolreprod57.2.335 (1997).

78 Isola, J. V. V. et al. Mild calorie restriction, but not 17alpha-estradiol, extends ovarian reserve and fertility in female mice. Exp Gerontol 159, 111669, doi:10.1016/j.exger.2021.111669 (2022).

79 Franasiak, J. M. et al. The nature of aneuploidy with increasing age of the female partner: a review of 15,169 consecutive trophectoderm biopsies evaluated with comprehensive chromosomal screening. Fertil Steril 101, 656–663 e651, doi:10.1016/j.fertnstert.2013.11.004 (2014).

80 Ben Yaakov, T., Wasserman, T., Aknin, E. & Savir, Y. Single-cell analysis of the aged ovarian immune system reveals a shift towards adaptive immunity and attenuated cell function. Elife 12, doi:10.7554/eLife.74915 (2023).

81 Masopust, D. & Soerens, A. G. Tissue-Resident T Cells and Other Resident Leukocytes. Annu Rev Immunol 37, 521–546, doi:10.1146/annurev-immunol-042617-053214 (2019).

82 Yuzen, D., Arck, P. C. & Thiele, K. Tissue-resident immunity in the female and male reproductive tract. Semin Immunopathol 44, 785–799, doi:10.1007/s00281-022-00934-8 (2022).

83 Wang, X. & Tian, Z. gammadelta T cells in liver diseases. Front Med 12, 262–268, doi:10.1007/s11684-017-0584-x (2018).

84 Hammerich, L. et al. Chemokine receptor CCR6-dependent accumulation of gammadelta T cells in injured liver restricts hepatic inflammation and fibrosis. Hepatology 59, 630–642, doi:10.1002/hep.26697 (2014).

85 Peng, X. et al. IL-17A produced by both gammadelta T and Th17 cells promotes renal fibrosis via RANTES-mediated leukocyte infiltration after renal obstruction. J Pathol 235, 79–89, doi:10.1002/path.4430 (2015).

86 Simonian, P. L. et al. gammadelta T cells protect against lung fibrosis via IL-22. J Exp Med 207, 2239–2253, doi:10.1084/jem.20100061 (2010).

87 Yan, X. et al. Deleterious effect of the IL-23/IL-17A axis and gammadeltaT cells on left ventricular remodeling after myocardial infarction. J Am Heart Assoc 1, e004408, doi:10.1161/JAHA.112.004408 (2012).

88 Bank, I. The Role of gammadelta T Cells in Fibrotic Diseases. Rambam Maimonides Med J 7, doi:10.5041/RMMJ.10256 (2016).

89 Zhang, M. & Zhang, S. T Cells in Fibrosis and Fibrotic Diseases. Front Immunol 11, 1142, doi:10.3389/fimmu.2020.01142 (2020).

90 Bruno, M. E. C. et al. Accumulation of gammadelta T cells in visceral fat with aging promotes chronic inflammation. Geroscience 44, 1761–1778, doi:10.1007/s11357-022-00572-w (2022).

91 Moutuou, M. M., Gauthier, S. D., Chen, N., Leboeuf, D. & Guimond, M. Studying Peripheral T Cell Homeostasis in Mice: A Concise Technical Review. Methods Mol Biol 2111, 267–283, doi:10.1007/978-1-0716-0266-9_21 (2020).

92 Nguyen, Q. P., Deng, T. Z., Witherden, D. A. & Goldrath, A. W. Origins of CD4(+) circulating and tissue-resident memory T-cells. Immunology 157, 3–12, doi:10.1111/imm.13059 (2019).

93 Wilkinson, P. C., Komai-Koma, M. & Newman, I. Locomotion and chemotaxis of lymphocytes. Autoimmunity 26, 55–72, doi:10.3109/08916939709009550 (1997).

94 Dong, Y. et al. The role of regulatory T cells in thymectomy-induced autoimmune ovarian disease. Am J Reprod Immunol 78, doi:10.1111/aji.12683 (2017).

95 Sharif, K. et al. Insights into the autoimmune aspect of premature ovarian insufficiency. Best Pract Res Clin Endocrinol Metab 33, 101323, doi:10.1016/j.beem.2019.101323 (2019).

96 Zhang, Z., Schlamp, F., Huang, L., Clark, H. & Brayboy, L. Inflammaging is associated with shifted macrophage ontogeny and polarization in the aging mouse ovary. Reproduction 159, 325–337, doi:10.1530/REP-19-0330 (2020).

97 Ahmadzadeh, K., Vanoppen, M., Rose, C. D., Matthys, P. & Wouters, C. H. Multinucleated Giant Cells: Current Insights in Phenotype, Biological Activities, and Mechanism of Formation. Front Cell Dev Biol 10, 873226, doi:10.3389/fcell.2022.873226 (2022).

98 Weivoda, M. M. & Bradley, E. W. Macrophages and Bone Remodeling. J Bone Miner Res 38, 359–369, doi:10.1002/jbmr.4773 (2023).

99 Kameda, T. et al. Estrogen inhibits bone resorption by directly inducing apoptosis of the bone-resorbing osteoclasts. J Exp Med 186, 489–495, doi:10.1084/jem.186.4.489 (1997).

100 Goretzlehner, G., Krause, B., Nehmzow, M. & Ulrich, U. [Endocrine diseases in pregnancy]. Z Arztl Fortbild (Jena) 84, 135–141 (1990).

101 Adamopoulos, I. E. et al. IL-17A gene transfer induces bone loss and epidermal hyperplasia associated with psoriatic arthritis. Ann Rheum Dis 74, 1284–1292, doi:10.1136/annrheumdis-2013-204782 (2015).

102 Anderson, J. M. Multinucleated giant cells. Curr Opin Hematol 7, 40–47, doi:10.1097/00062752-200001000-00008 (2000).

103 Fais, S. et al. Multinucleated giant cells generation induced by interferon-gamma. Changes in the expression and distribution of the intercellular adhesion molecule-1 during macrophages fusion and multinucleated giant cell formation. Lab Invest 71, 737–744 (1994).

104 Lind, A. K. et al. Collagens in the human ovary and their changes in the perifollicular stroma during ovulation. Acta Obstet Gynecol Scand 85, 1476–1484, doi:10.1080/00016340601033741 (2006).

105 Liu, G. Y. & Sabatini, D. M. mTOR at the nexus of nutrition, growth, ageing and disease. Nat Rev Mol Cell Biol 21, 183–203, doi:10.1038/s41580-019-0199-y (2020).

106 Schneider, A. et al. The Interconnections Between Somatic and Ovarian Aging in Murine Models. J Gerontol A Biol Sci Med Sci 76, 1579–1586, doi:10.1093/gerona/glaa258 (2021).

107 Young, M. D. & Behjati, S. SoupX removes ambient RNA contamination from droplet-based single-cell RNA sequencing data. Gigascience 9, doi:10.1093/gigascience/giaa151 (2020).

108 Satija, R., Farrell, J. A., Gennert, D., Schier, A. F. & Regev, A. Spatial reconstruction of single-cell gene expression data. Nat Biotechnol 33, 495–502, doi:10.1038/nbt.3192 (2015).

109 Liu, Y. et al. Single-Cell Profiling Reveals Divergent, Globally Patterned Immune Responses in Murine Skin Inflammation. iScience 23, 101582, doi:10.1016/j.isci.2020.101582 (2020).

110 Nguyen, H. T. T., Guevarra, R. B., Magez, S. & Radwanska, M. Single-cell transcriptome profiling and the use of AID deficient mice reveal that B cell activation combined with antibody class switch recombination and somatic hypermutation do not benefit the control of experimental trypanosomosis. PLoS Pathog 17, e1010026, doi:10.1371/journal.ppat.1010026 (2021).

111 McGinnis, C. S., Murrow, L. M. & Gartner, Z. J. DoubletFinder: Doublet Detection in Single-Cell RNA Sequencing Data Using Artificial Nearest Neighbors. Cell Syst 8, 329–337 e324, doi:10.1016/j.cels.2019.03.003 (2019).

112 Jin, S. et al. Inference and analysis of cell-cell communication using CellChat. Nat Commun 12, 1088, doi:10.1038/s41467-021-21246-9 (2021).

113 Isola, J. V. V. et al. 17alpha-Estradiol promotes ovarian aging in growth hormone receptor knockout mice, but not wild-type littermates. Exp Gerontol 129, 110769, doi:10.1016/j.exger.2019.110769 (2020).

114 Saccon, T. D. et al. Primordial follicle reserve, DNA damage and macrophage infiltration in the ovaries of the long-living Ames dwarf mice. Exp Gerontol 132, 110851, doi:10.1016/j.exger.2020.110851 (2020).

115 Anderson, K. G. et al. Intravascular staining for discrimination of vascular and tissue leukocytes. Nat Protoc 9, 209–222, doi:10.1038/nprot.2014.005 (2014).

116 Ruxton, G. D. & Neuhäuser, M. When should we use one-tailed hypothesis testing? 1, 114–117, 10.1111/j.2041-210X.2010.00014.x (2010).

